# An interpretable neural network unveils higher-order epistasis in large protein sequence-function relationships

**DOI:** 10.1101/2024.09.22.614318

**Authors:** Palash Sethi, Juannan Zhou

**Affiliations:** Department of Biology, University of Florida, Gainesville, FL, 32611

## Abstract

Protein sequence–function relationships are inherently complex, as amino acids at different positions can interact in highly unpredictable ways. A key question for protein evolution and engineering is how often epistasis extends beyond pairwise interactions to involve three or more positions. Although experimental data has accumulated rapidly in recent years, addressing this question remains challenging, as the number of possible interactions is typically enormous even for proteins of moderate size. Here, we introduce an interpretable machine learning framework for studying higher-order epistasis scalable to full-length proteins. Our model builds on the transformer architecture, with key modifications allowing us to assess the importance of higher-order interactions by fitting a series of models with increasing complexity. Applying our method to 10 large protein sequence-function datasets, we found that while additive effects explain the majority of the variance, within the epistatic component, the contribution of higher-order epistasis ranges from negligible to up to 60%. We also found higher-order epistasis is particularly important for generalizing locally sampled fitness data to distant regions of sequence space and for modeling an additional multi-peak fitness landscape. Our findings suggest that higher-order epistasis can play important roles in protein sequence-function relationships, and thus should be properly considered in protein engineering and evolutionary data analysis.

## Introduction

Understanding how amino acid sequence determines protein function is critical for biomedical engineering, synthetic biology, and evolutionary biology [1–4]. High-throughput techniques such as deep mutational scanning (DMS) have enabled the functional characterization of thousands to millions of protein variants in a massively parallel manner [5–24]. A general observation from these empirical studies is that while sequence–function relationships can often be reasonably approximated by the independent contributions of mutations, epistatic interactions between amino acids also play important roles and must be properly accounted for to accurately model protein function [25–29].

Although pairwise interactions between amino acids are both predicted by biophysical models and widely supported by empirical data [30–34], our understanding of how frequently epistasis extends to involve three or more positions remains incomplete. As these higher-order interactions can pose significant conceptual and practical challenges for deciphering protein sequence–function relationships, a thorough assessment of their prevalence is essential for informing future experimental designs and the development of analytical methods [4, 35]. Current evidence on this question is mixed: some studies suggest that higher-order epistasis is widespread and plays a significant role in protein function [28, 30, 36–38], while others argue that protein sequence–function landscapes are simpler and epistasis beyond pairwise interactions is rare [31, 33].

For example, a recent paper [31] investigated the genetic architecture for 20 protein datasets derived from combinatorial mutagenesis by comparing the predictive performance of pairwise and higher-order epistatic models, while simultaneously fitting a sigmoidal function to account for non-specific epistasis. In contrast to conclusions from some of the earlier studies, the authors observed small to negligible contribution of higherorder epistasis and argue that most protein fitness landscapes can be well characterized by only additive effects and pairwise interactions. However, a key limitation of this study is that it is based on small-scale protein sequence-function relationships, involving either exhaustive amino acid substitutions at no more than four sites or simultaneous single amino acid substitutions at up to 20 sites.

To draw general conclusions about the role of higher-order epistasis, we must extend these tests to larger protein sequence-function relationships as interactions beyond pairwise epistasis become more probable due to simple combinatorics and biophysical principles. Historically, researchers have quantified the contribution of higher-order epistasis by decomposing complete fitness landscapes into orthogonal interaction coefficients [28, 39]. These methods are only applicable to small sequence spaces as they require (near) combinatorially complete datasets. When dataset is incomplete, researchers typically resort to fitting parametric regressions and assessing the contribution of epistasis by comparing the prediction performance on test datasets for models with and without certain epistatic components [23, 31, 40]. However, naive implementation of parametric regression methods to model full-length proteins leads to extremely high computational demands, overfitting, and lack of model interpretability due to the exponential growth of model parameters.

Although nonparametric methods represent a desirable, and highly scalable approach by modeling higherorder interactions without explicitly fitting regression coefficients [29, 41–46], generalizing it to account for the confounding effects of non-specific epistasis using a global nonlinearity [31, 47, 48] is not straightforward due to the resulting non-Gaussian likelihood [49].

Artificial neural networks (ANNs) have recently been widely applied to model sequence-function relationships [19, 50, 51]. Their flexibility makes them well-suited to capturing complex, higher-order epistatic interactions. However, naïvely applying ANNs provides limited insight into the prevalence or structure of higher-order epistasis, as there is no direct correspondence between the architecture of a neural network and the structure of epistasis it models. Notably, there have been active development of interpretable deep learning models for studying the structure of epistasis based on protein sequence-function data. For instance, network architectures accounting for the joint effect of global epsitasis and independent contributions of mutations have been widely adapted for its model simplicity and intuitive behavior [11, 47, 52, 53]. This class of models have also been generalized to explicitly model pairwise or even higher-order epistasis[54, 55]. Furthermore, advances in interpretable AI, such as Shapley values and Shapley interaction indices [56, 57], can potentially be adapted for interpreting black-box ANNs to estimate the marginal effects of positions and interactions among pairs or subsets of loci. Although promising, these methods are still not exempt from the exponential scaling of their model complexity with interaction order faced by conventional regression methods.

To address this challenge, we developed a modified transformer architecture. The key feature of our method is that it allows us to explicit control the maximum order of epistasis the network fits by simply adjusting the number of attention layers. This design allows us to systematically assess the contribution of higher-order interactions by fitting a series of models with increasing epistatic complexity and comparing their predictive performance. Crucially, unlike traditional regression-based approaches, our method captures higher-order interactions implicitly through learned neural network weights. As a result, model complexity does not grow exponentially with sequence length or interaction order. This enables us to model specific higher-order epistasis, while accounting for global epistasis in DMS datasets for full-length protein, whereas previous methods can only achieve this for a limited number of mutagenized positions.

We begin by introducing the core ideas of our method and demonstrating its desirable behavior on a simulated dataset. We next present our findings on the importance of higher-order epistasis across 10 largescale combinatorial protein DMS datasets. We also demonstrate how higher-order epistasis contributes to out-of-distribution predictions and determines the structure of a multi-peak fitness landscape comprising four GFP orthologs. Finally, we present an exploratory analysis of a trained neural network to gain insights into the architecture of higher-order interactions in the protein GRB2-SH3.

## Results

### Epistatic transformer for fitting higher-order epistasis

We aim to quantify the contribution of specific higher-order epistasis to the total phenotypic variance measured for empirical protein sequence-function relationships. Our overall strategy is to fit models of increasing complexity to these datasets and examine how inclusion of higher-order interactions improves model generalization to held-out test genotypes.

To account for epistasis resulting from interactions among specific mutations, as well as from nonlinear scaling on the observation scale (i.e., nonspecific or global epistasis[3, 47, 48]), we study models of the general form

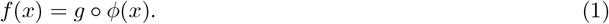

Here *x* = *x*_1_, *x*_2_, …, *x*_*L*_ is the amino acid sequence of a protein of length *L. ϕ*(*x*) is a function of the independent effects of individual amino acids as well as specific epistatic interactions, and can be expressed as a one-hot model [58]

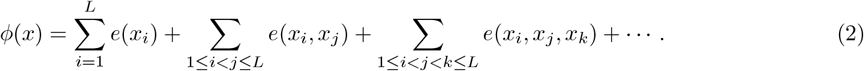

Specifically, *e*(*x*_*i*_) captures the independent effect of residue *x*_*i*_ at site *i, e*(*x*_*i*_, *x*_*j*_) captures the interaction effect between residues *x*_*i*_ and *x*_*j*_. *e*(*x*_*i*_, *x*_*j*_, *x*_*k*_) accounts for three-way interaction, etc. In our framework, *ϕ* may include epistatic interactions among up to *K* ≤*L* positions. *g* is a nonlinear monotonic function that transforms the latent phenotype *ϕ*(*x*) to the measurement scale for modeling global epistasis [47, 51–54]. Our strategy is to fit a series of models of the form Eq. 1 by varying the complexity of the function *ϕ* with increasing values of *K*, in order to examine if higher *K* leads to better generalizability of the trained model to novel genotypes.

Fitting the function *ϕ* is challenging when higher-order interactions are considered, as the number of regression coefficients in Eq. 2 grows roughly exponentially with interaction order. For example, for a protein with 100 mutagenized positions, there are 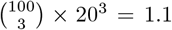 billion three-way interactions and 511 billion four-way interactions. Thus for large protein sequence spaces, current methods that directly fit the regression coefficients in Eq. 2 can at most capture interactions for pairs of sites [54, 55].

Neural networks offer an attractive alternative due to their ability to capture complex interactions using neural network weights, instead of explicitly fitting regression coefficients. It thus could potentially avoid the exponential increase in model complexity with interaction order. However, standard neural network architectures inherently induce interactions of all orders through the numerous nonlinear functions in their hidden layers, making it impossible to decompose the model in the form of Eqs. 1 and 2 for testing if incorporating specific pairwise or higher-order epistasis improves model performance.

Here, we developed a novel neural network architecture, the epistatic transformer, for modeling specific epistatic interactions of fixed orders. The model allows one to fit Eq. 1, with *ϕ* containing specific epistasis among up to *K* = 2^*M*^ sites with *M* attention layers, such that by setting *M* = 1, 2, 3, …, we can fit pairwise, four-way, eight-way, or higher-order interaction models. The high-level model architecture is shown in Figure 1a. The input protein sequence *x*_1_, …, *x*_*L*_ is first used to generate amino acid embeddings 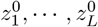 of dimension *d*. The embeddings are then passed through *M* modified multi-head attention (MHA) layers [59], such that the output embeddings 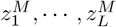 contain specific interactions exactly up to order 2^*M*^ . We then calculate a single weighted sum of the flattened embeddings, which is passed through a nonlinear function *g* for modeling global epistasis.

**Figure 1.**
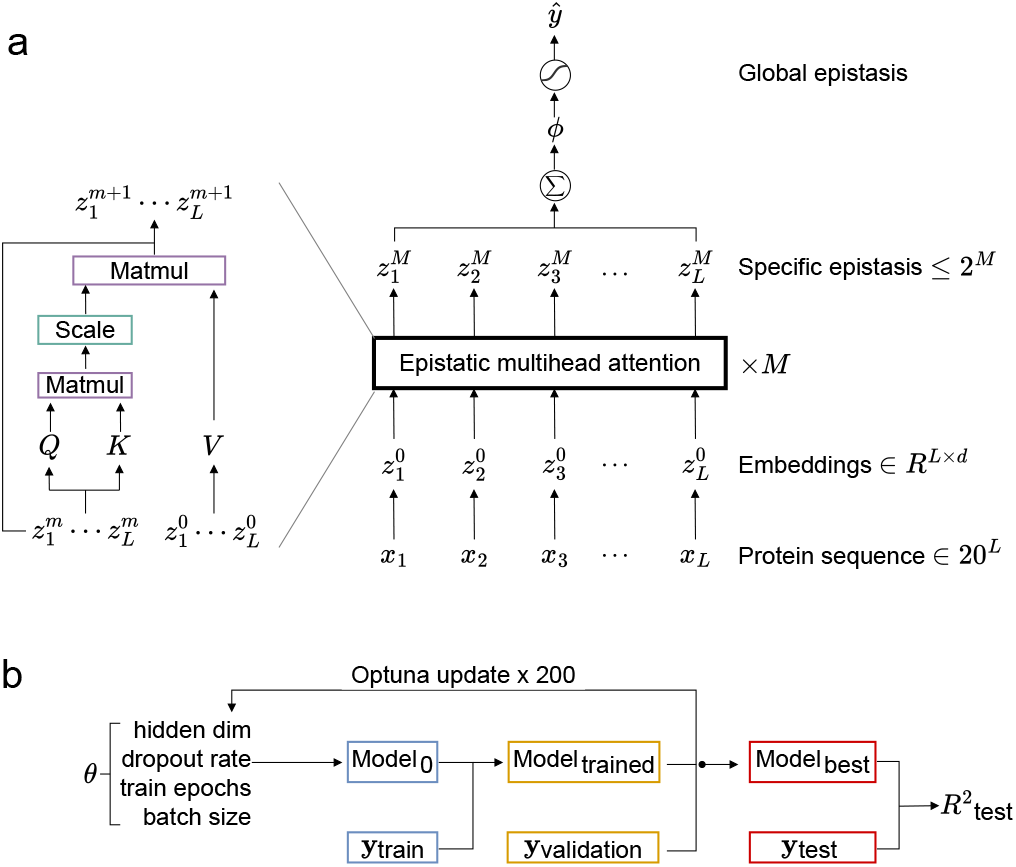
Epistatic transformer for jointly modeling fixed-order specific epistasis and nonspecific epistasis. a. Overall model architecture. The input amino acid token *x*_*l*_ is first used to generate position-specific amino acid embeddings 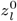. The embeddings are then passed through *M* layers of epistatic multi-head attention (MHA) such that the output embeddings contain specific epistasis up to order 2^*M*^ . Left: Details for a single epistatic MHA layer. Hidden states *z*^*m*^ from the previous layer is passed through linear layers to generate the query (*Q*) and key (*K*) tensors, while the raw embeddings *Z*^0^ are used directly as the value (*V* ) tensor. Attention weights are calculated by taking scaled dot products between *Q* and *K*, which are used to generate the final output of this layer. This bypassing of the raw embedding *Z*^0^, together with the removal of LayerNorm, softmax operation on the attention weights, and the feedforward layer, allow us to model only interactions among up to 2^*M*^ sites when using *M* epistatic MHA layers. b. Automated hyperparameter search scheme using Optuna [60]

In a standard attention layer, input embeddings are linearly projected to form query (*Q*), key (*K*), and value (*V* ) matrices. For a focal position *i*, attention scores are computed as the scaled dot product between its query vector *q*_*i*_ and all keys, 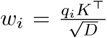. The output at position *i* is then obtained as a weighted sum of values, softmax(*w*_*i*_)*V* . Intuitively, a single attention layer induces pairwise interactions. And stacking additional layers enables the model to capture interactions among interactions learned in earlier layers. As a result, the receptive field of a position and the effective order of interactions grow roughly exponentially with depth. This property makes it possible to probe the importance of different orders of epistasis by fitting a series of models with increasing numbers of attention layers.

However, generic transformer architecture is still not suitable for fitting specific epistasis, as standard MHA layers have the same problem of mixing global epistasis and specific epistasis operations as other neural network models. Here we made three key modifications to the MHA layer (Figure 1a, left panel; Methods) to allow us to precisely control the orders of specific epistasis allowed in the latent phenotype *ϕ* and to better fit the idiosyncratic epistatic interactions in the data.

First, we remove all nonlinear operations in the hidden layers of the model, including the softmax function applied to the attention weights, LayerNorm [61], and the final feedforward network within each transformer encoder layer. This modification ensures that the *ϕ* function we infer only contains specific interactions. Second, we replace the value (*V* ) input to each MHA layer with the raw amino acid embedding 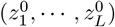. This key modification ensures that the order of specific epistasis in our model scales exactly as 2^*M*^ with the number of layers *M* (see Supplemental Methods for proof). Thus, by setting *M* = 1, 2, 3, we are able to fit Eq. 2 with up to pairwise, 4th-order, and 8th-order interactions, respectively. Third, we use a special positional encoding for the input amino acids, where the original amino acid token *x*_*l*_ is transformed to 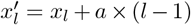 (*a* is the alphabet size), effectively allowing us to treat amino acids at different positions in completely different ways. Our experiments show that this is more efficient than conventional positional embeddings in capturing the range of interactions between residues in real protein datasets.

To model global epistasis, we use a nonlinear function *g* to map the weighted sum of the flattened embeddings from the final epistatic MHA layer to the model prediction. Specifically, we evaluated a flexible family of function in which *g* is the sum of varying numbers of independent sigmoid functions [19, 54]. We found that a single scaled sigmoidal function with four parameters, 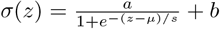, performs as well as complex functions with multiple sigmoid functions on the empirical datasets (Supplemental Figure 1). Thus, we use this activation function to model nonspecific epistasis in all subsequent analyses.

Preliminary experiments showed that the performance of epistatic transformer is sensitive to choices of model hyperparameters. Thus, to ensure we always fit the best model with *M* epistatic MHA layers to a given dataset, such that improvement over the model of *M* − 1 layers is solely due to inclusion of higher-order epistasis, we used Optuna [60] to optimize a number of hyperparameters on an independent validation dataset (Figure 1b). The model trained under the optimal hyperparameters was evaluated on the test dataset to produce the *R*^2^ value.

### Epistatic transformer captures specific higher-order epistasis in a simulated fitness landscape

We first apply our model to a simulated dataset to characterize its behavior when the ground truth is known. We simulated a sequence-function map for a binary sequence space with 13 sites, with specific epistatic interactions up to order 8 (see Supplemental Methods). The choice of a small genotype space allows us to easily generate data for all possible genotypes and precisely control the formal structure of epistasis in the simulated fitness landscape, which is not tractable in larger sequence spaces.

Figure 2a summarizes the structure of epistasis in our simulated data in terms of the fraction of total phenotypic variance explained by additive effects and epistatic interactions up to a given order (i.e., cumulative variance explained). Specifically, additive effects alone account for 10% of the total variance, while additive and pairwise interactions together explain 30%. Furthermore, 70% of the phenotypic variance arises from interactions involving up to four sites, and considering interactions among up to eight sites explains all variance.

**Figure 2.**
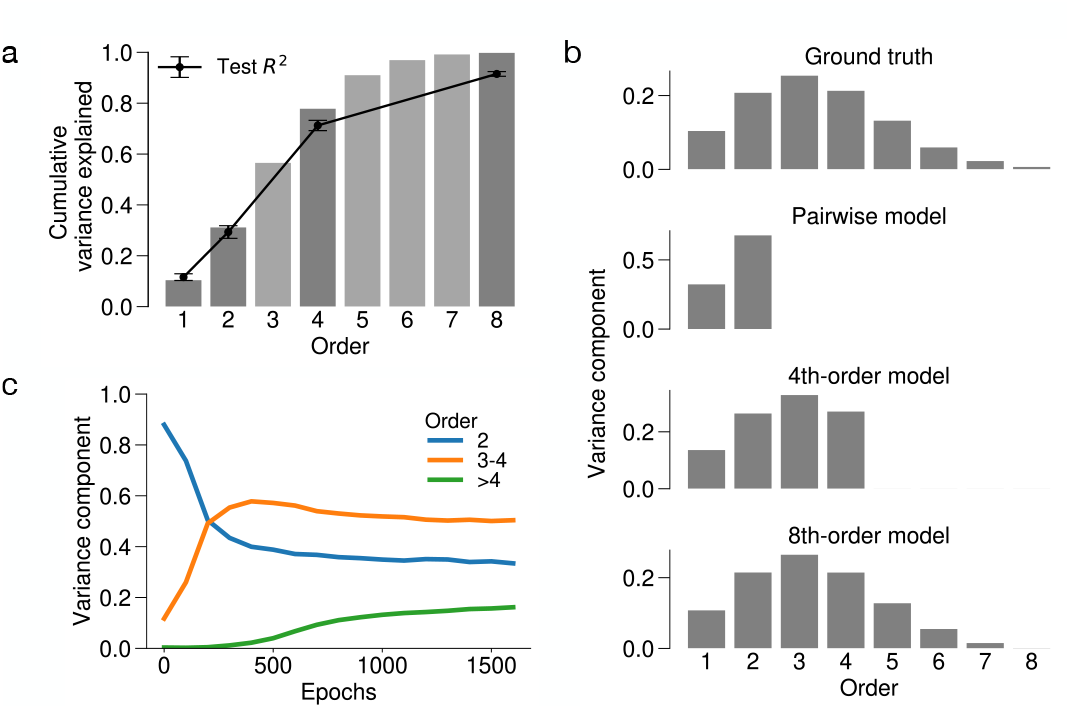
Epistatic transformer is able to capture the genetic architecture of simulated sequence-function relationships. Data was simulated for a binary fitness landscape with 13 sites, with specific epistasis among up to 8 sites. a. Test *R*^2^ can be used to estimate cumulative variance components. Bar plot shows the cumulative variance for each order of interaction (proportion of variance due to epistasis up to a given order). Error plot shows the test *R*^2^ for additive, pairwise, four-way, and eight-way interaction models using epistatic transformer to 90% of simulated data. Error bars correspond to one standard deviation with 5 random train-test splits. b. Epistatic transformer recapitulates true variance components. Bar plots correspond to variance components(proportion of variance explained by interactions of a given order) for the simulated landscape, the pairwise, 4th-order, and 8th-order models. Note that pairwise model only contains interactions up to second order, and 4th-order model contains interactions among up to 4 positions, while 8th-order model model contains interactions among up to 8 positions with variance components aligning with the ground truth. c. Epistatic transformer model becomes increasingly complex with longer training. The three curves represent the variance component decomposition of the full landscape inferred by the 8th-order model at each training epoch. Blue: variance components for additive and pairwise interactions. Orange: variance components for 3rd-order and 4th-order interactions. Green: variance components for interaction order *>* 4.

Next, we examine if we can infer the structure of specific epistasis in the data by training a series of epistatic transformer models to predict the phenotype *ϕ*. Specifically, we fit epistatic transformer to 90% of randomly sampled data and examine its out-of-sample performance when the number of epistatic MHA layer is set to *M* = 1, 2, or 3, corresponding to pairwise, 4th-order, and 8th-order interaction models due to the 2^*M*^ scaling relationship. We see that the test *R*^2^ increases with the maximum interaction order of the model and remains slightly below—but close to—the ground truth cumulative variance explained. These results confirm that our model can be used to provide lower bounds for the contribution of pairwise and higher-order epistasis when fitted to randomly sample training data.

Figure 2b shows the variance components (the fraction of variance explained by additive effects or epistasis of specific orders) of the simulated data and the predictions made by the low-order (*M* = 1, 2) and all-order epistatic transformer models (*M* = 3). We see that for the low-order interaction models, the variance components are truncated at the highest order the model can theoretically capture. For instance, a 4th-order interaction model with 2 epistatic MHA layers only contains interactions among up to 4 sites. In contrast, the variance components of the 3-layer model contains all 8 orders of interactions, and faithfully recapitulate the ground-true variance components (Figure 2b, top panel). Together, this result confirms the 2^*M*^ scaling relationship we proved mathematically, and that the all-order epistatic transformer model can capture arbitrary structures of specific epistasis.

Lastly, we examine how the model learns higher-order interactions over the course of training. In Figure 2c, we plot the variance components of the 8th-order model attributable to additive and pairwise interactions, 3rd + 4th-order interactions, and interactions of order *>* 4 at different training epochs. Early in training, the model primarily captures additive and pairwise components (blue curve). As training progresses, higher-order interactions become increasingly prominent (orange and green curves), leading to convergence of the reconstructed variance components to the ground truth. This interesting behavior suggests that the model first learns simpler features of the sequence–function relationship before progressively capturing more complex interactions.

### Application to experimental protein sequence-function data

To characterize the importance of higher-order interactions in empirical protein fitness landscapes using the epistatic transformer model, we curated a dataset derived from 10 combinatorial deep mutational scanning (DMS) experiments. To ensure sufficient training data for the model to detect higher-order interactions, we selected datasets with a large number of mutated positions and a high proportion of genotypes more than three mutations away from the reference wild type sequence (Methods).

The resulting 10 datasets (Table 1) encompass a diverse set of proteins and phenotypes including protein nabundance, binding, fluorescence, and cellular fitness. Importantly, the selected experiments were conducted in sequence spaces significantly larger than datasets used in previous studies, with mutations spanning tens of amino acid sites, extending up to the entire length of the protein (e.g. the GFP datasets). The chosen datasets also exhibit a wide range of data distribution in terms of the of mutational distance to the wild type and the sparsity of the data (Supplemental Figure 2).

**Table 1:**
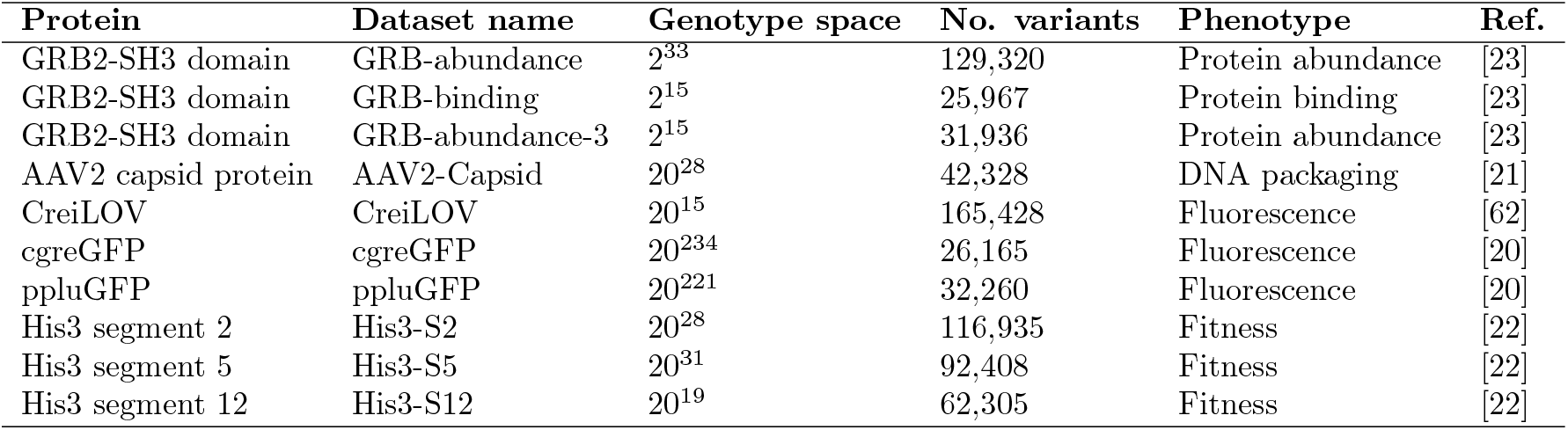
Combinatorial protein mutagenesis datasets used in this paper.

For each dataset, we assigned random subsets of all measured genotypes as training data. We used epistatic transformer models to fit pairwise, 4th-order, and 8th-order models in the form of Eq. 1, corresponding to models with *M* = 1, 2, and 3 epistatic MHA layers. In addition, we also fit an additive model with a sigmoid activation as a performance baseline. To ensure that we fit the best model to the training data, we used the procedure in Figure 1b to identify the optimal hyperparameter combination using cross validation with 200 tuning steps (Methods).

We first observe that the baseline additive model with a global epistasis activation often provides a reasonably good fit to the data and explains a substantial fraction of the total variance (test *R*^2^ *>* 0.55 for all datasets when trained on 80% of randomly sampled genotypes; Supplemental Figure 3). For several datasets, the additive model explains up to 90% of the variance in the test data (e.g., GRB-abundance-2, His-S5, and His-S12). We also found that incorporating global epistasis uniformly improves over the purely additive model (Supplemental Figure 3), with gains in test *R*^2^ ranging from less than 10% for the three GRB datasets to as much as 30% for the cgreGFP dataset.

Having accounted for the contributions of additive effects and global epistasis, we next characterize the structure of specific epistasis using two metrics. First, we use the difference in test *R*^2^ between the most complex model (3 epistatic MHA layers) and the additive model, 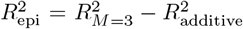, to measure the joint contribution of all orders of specific epistasis in the data.

**Table 2:** Formulas for calculating the percent epistatic variance for different orders of specific epistatic interactions. 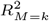 is the test *R*^2^ value for an epistatic transformer model with *k* epistatic MHA layers. 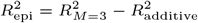.

| Orders of epistasis | % epistatic variance |
| --- | --- |
| 2 | $(R_{M=1}^2 - R_{\text{additive}}^2) / R_{\text{epi}}^2$ |
| 3, 4 | $(R_{M=2}^2 - R_{M=1}^2) / R_{\text{epi}}^2$ |
| 5, 6, 7, 8 | $(R_{M=3}^2 - R_{M=2}^2) / R_{\text{epi}}^2$ |

Next, to quantify the contribution of specific orders of epistasis, we exploited the nested structure of our modeling framework by comparing the test *R*^2^ of model with *M* epistatic MHA layers and the model with *M* − 1 layers. For example, a model with 2 epistatic MHA layers captures 3rd- and 4th-order epistatic interactions in addition to the pairwise interactions modeled with a single attention layer. The difference in their test *R*^2^ values therefore corresponds to the variance attributable to higher-order epistatic effects uniquely captured by the higher-order model. We further normalize this difference by 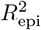 to define the ‘percent epistatic variance’ for comparing of the relative contributions of epistasis across different orders (Table 2).

We first observe that the contribution of specific epistasis can vary substantially across datasets, with 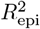 (top right corner of each panel in Figure 3) ranging from 0.01 (His3-S5 and His3-S12) to 0.23 (GRB-abundance). Second, we found that while pairwise interactions are typically the predominant form of epistasis, consistent with previous findings, higher-order epistasis can also play moderate to substantial roles. We found the strongest contribution of higher-order epistasis in the GRB-abundance dataset, wherein three-way and four-way interactions together account for 62% of the epistatic variance (14.2% of total variance), compared to 36% attributed to pairwise interactions (Figure 3). In contrast, we observed that interactions of order *>* 4 account for small amount of epistatic variance, suggesting that three-way and four-way interactions are the predominant forms of epistasis in this dataset.

**Figure 3.**
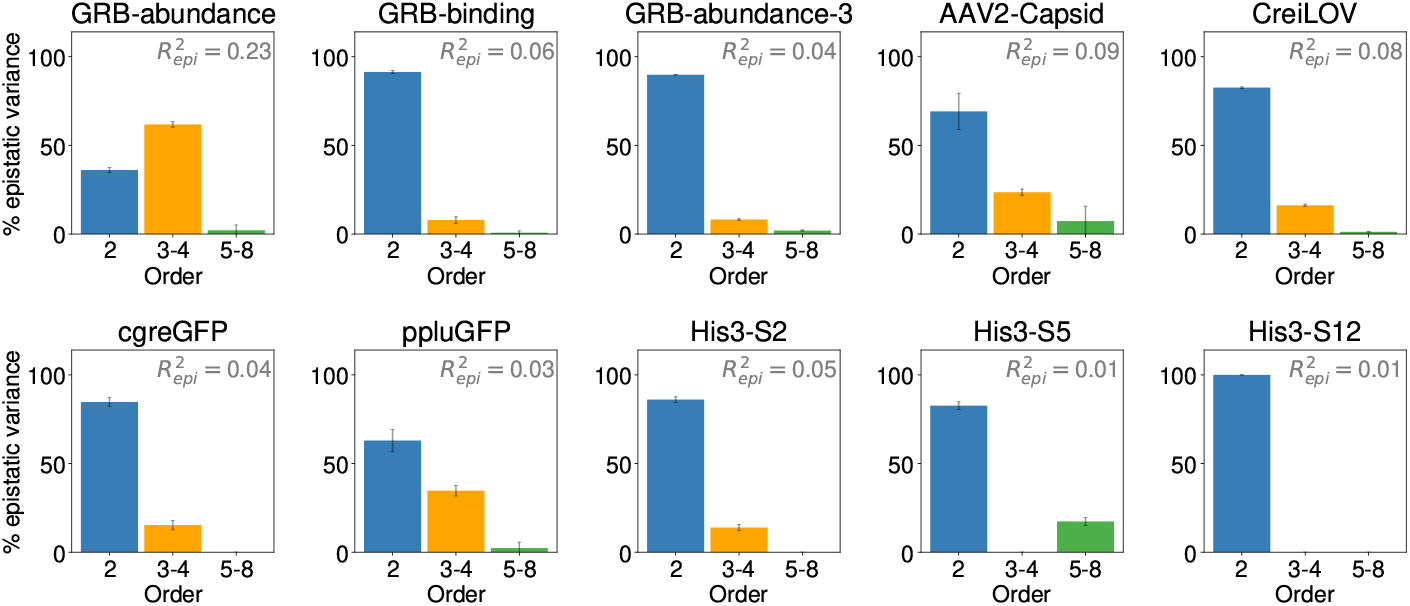
Importance of pairwise and higher-order specific epistasis in 10 experimental protein-sequence function datasets. For each dataset, pairwise, 4th-order and 8th-order models were fit using epistatic transformer with 1, 2, and 3 layers of epistatic MHA, along with an additive model. All models contain a final sigmoid activation function mapping a scalar value to the measurement scale for modeling non-specific epistasis. Models were fit to 80% of training data generated by randomly sampling all available data and evaluated on random test genotypes. In each panel, the number 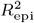 on the upper right corner is equal to the proportion of total variance in the test data explained by all orders of specific epistasis, equal to the difference between the *R*^2^ of the 8th-order epistatic transformer model and the additive model. Importance of epistatic interactions of different orders is measured by percent epistatic variance, equal to the gain in *R*^2^ by fitting an additional layer of epistatic MHA, normalized by 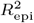. For example, the percent epistatic variance due to pairwise interactions is equal to the difference in *R*^2^ between the pairwise and additive model divided by 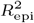, and the percent epistatic variance due to 3-way and 4-way interactions is equal to the difference in *R*^2^ by the 4th-order model and the pairwise divided by 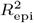 (Table 2). Error bars represent 1 standard deviation calculated from 3 replicates.

We also observed moderate contributions by higher-order epistasis in other datasets (e.g. AAV2-Capsid, ppluGFP, and CreiLOV), where epistasis of order 3-8 collectively accounts for roughly 1*/*5 to 1*/*3 of the total epistatic variance. Interestingly, we also found that higher-order epistasis can make more substantial contributions when training data is more sparse, sometimes accounting for *>* 80% of the total epistatic variance (e.g. the AA2-Capisd and cgreGFP dataset, Supplemental Figure 4). Similar phenomena have been observed previously, for instance, in non-parametric kernel methods [29, 41]. This suggests that higher-order models may, counterintuitively, provide a more parsimonious fit to sparse training data than the pairwise interaction model. Finally, we compare the performance between an epistatic transformer with 3 layers of epistatic MHA and a transformer model with generic attention mechanism. We observe that generic MHA yields at most a 1% improvement in *R*^2^ (Supplemental Figure 5). This suggests that the epistatic transformer incurs minimal loss in accuracy in exchange for substantially improved interpretability.

### Improvement in prediction accuracy in higher-order models is due to specific epistasis

In order to understand how higher-order epistasis improves model prediction, we next examine the residual structure of the pairwise and higher-order models fit to AAV2-Capsid and GRB-abundance, the two datasets with the largest contribution of higher-order interactions. In Figure 4, we show scatter plots comparing the actual experimental measurements *y* of the test genotypes vs. (1) the predicted latent phenotype *ϕ* (left panel for each dataset), and (2) the final predicted phenotype *ŷ*, after activation by the global epistasis function *g* (right panels). We first see that both datasets contain a prominent nonlinear relationship (red curve on the left panels), where genotypes with low and high predicted *ϕ* values are bounded by the sigmoid function. Next, we see that the final predictions for both models and datasets appear to have a linear correlation with the measurements (right panels). We also notice that the nonlinear functions for the pairwise and 8th-order models are highly similar, although the nonlinear function in these two models are fit separately.

**Figure 4.**
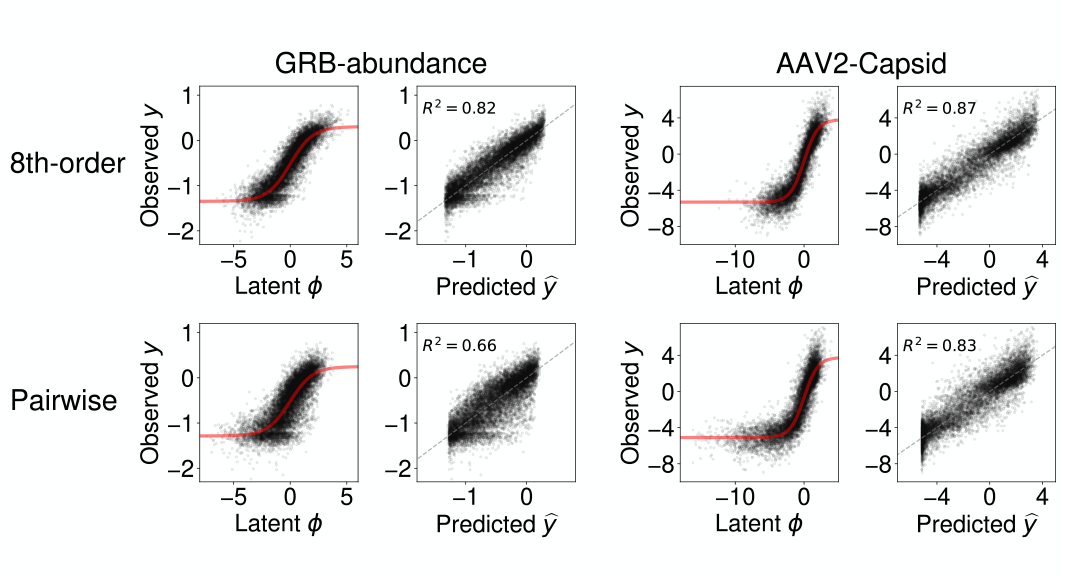
Improvement in prediction accuracy in higher-order models is due to their abilities to fit specific epistasis. Scatter plots show the observed phenotypes (*y*) vs. latent model predictions (*ϕ*), or the final model predictions 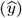) for test genotypes, for the pairwise and 8th-order epistatic transformer models. Models were fit to 80% of randomly sampled genotypes in the GRB-abundance and AAV2-Capsid datasets.

Together, these results confirm that both the pairwise and higher-order epistatic transformer models have adequately captured the non-specific epistasis in the data, evident by the lack of a nonlinear relationship between *y* and *ŷ*. This suggests that the improvement by the higher-order order epistatic model over the pairwise model is due to its ability to fit higher-order epistasis in the residual variance.

It is noteworthy that, for both the GRB-abundance and AAV2-Capsid datasets, the 8th-order model offers better predictions across the entire range of measured phenotypes. This is evident from the reduced spread of test data points for different observed *y* values, both on the latent and predicted phenotype scales. Moreover, the improvement in prediction accuracy is most significant for genotypes with high or low observed *y* values (Supplemental Figure 6), indicating the higher-order model’s ability to provide more parsimonious fits for genotypes at the extremes of the measurement scale.

### Higher-order epistasis is important for out-of-distribution predictions

Many datasets analyzed so far consist of genotypes generated by mutating a single wild-type or reference sequence, which inevitably results in highly localized samples within the whole genotype space. It is therefore critical to evaluate whether models trained on these local data generalize to genotypes distant from the training set. In this section, we characterize the predictive performance of epistatic transformer models for out-of-distribution genotypes. We focus on the AAV2-Capsid and cgreGFP datasets, which include dense measurements near the wild type and sparse higher-order mutants at larger mutational distances (Supplemental Figure 2).

In the previous section, we found incorporating higher-order epistasis leads to moderate improvement in predictive performance on randomly sampled test genotypes for both datasets. Here, we are interested in the impact of higher-order epistasis on model prediction for genotypes at varying distances to the data. Specifically, we use the mean Hamming distance 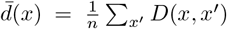 to measure the proximity of a genotype *x* to all genotypes *x*^*′*^ in the training data. We then stratified test genotypes into discrete bins by their mean Hamming distance and quantify the contribution of higher-order epistasis for each distance class by calculating the percent epistatic variance (Table 2) using models trained on 20% of randomly chosen genotypes.

For the AAV2-Capsid dataset, we first observed that the density of test genotypes decreases with mean distances, as a result of the localized data distribution (Figure 5a, top panel). This is accompanied by an overall decrease in the observed phenotypic scores (*y*) for genotypes in each distance class (Figure 5a, second panel). Furthermore, we also found that model predictive performance decreases monotonically with mean distance, such that the *R*^2^ of the additive model starts out at around 0.8 for nearby genotypes and drops to 0.4 for distant genotypes (Figure 5a, third panel). The *R*^2^ for the 8th-order epistatic transformer model with 3 epistatic MHA layers showed similar trend of decline with increasing distance, but maintained a relatively constant advantage over the additive model, with 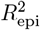 ranging from 0.05 to 0.14.

**Figure 5.**
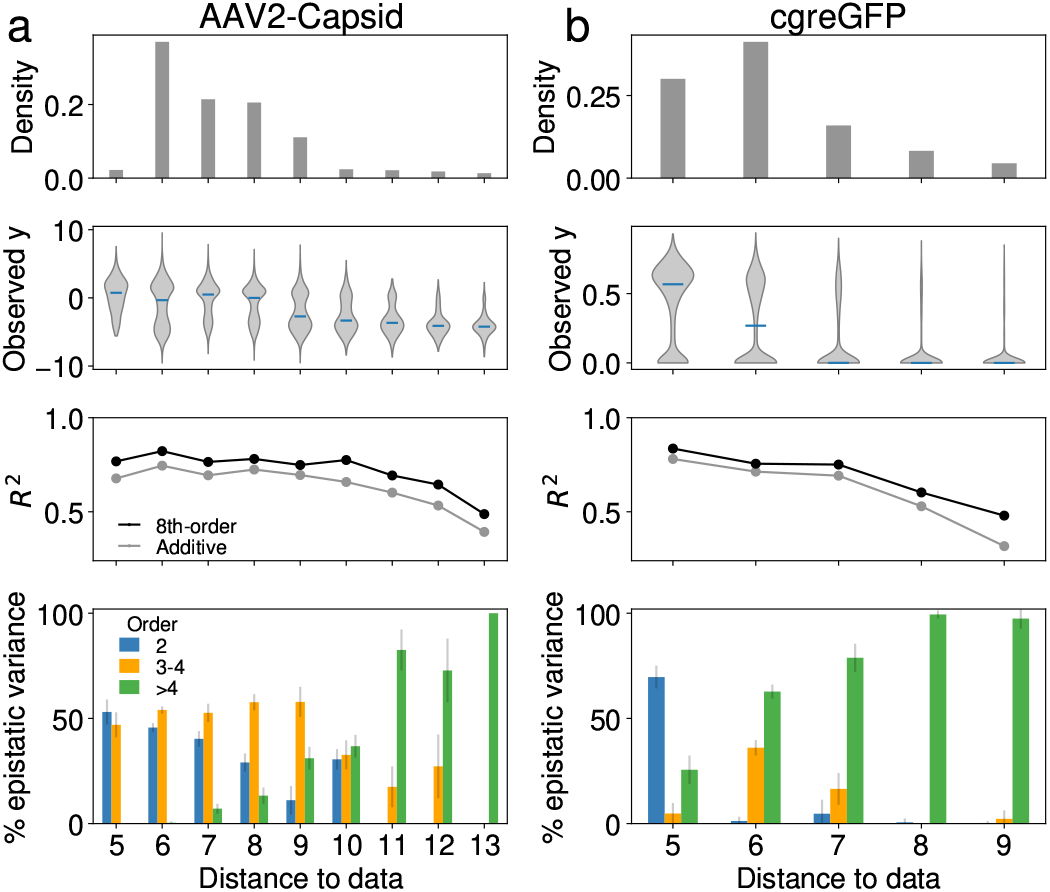
Importance of higher-order epistasis in predicting phenotypes for distant genotypes in the AAV2-Capsid (a) and the cgreGFP (b) datasets. For each dataset, genotypes are binned to discrete distance classes by their mean Hamming distances to the training data comprising 20% of randomly sampled genotypes. For both datasets, we retain only distance classes where the additive model has a test *R*^2^ *>* 0.3. Top panel: Distribution of mean Hamming distance in the randomly sampled test data. Second panel: Distribution of the observed phenotypic values (*y*) for each distance class. Third panel: Test *R*^2^ under the additive model and the 8th-order epistatic transformer for genotypes at different distance classes. The gap between the two curves is equal to 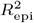 for each distance class. Bottom panel: Importance of specific pairwise and higher-order epistasis at different distance classes, measured by percent epistatic variance, equal to the gain in *R*^2^ by fitting an additional layer of epistatic MHA, normalized by 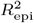 (Table 2). All metrics were calculated for models fit to one training sample consisting of 20% of randomly sampled genotypes. Error bars represent 1 standard deviation calculated by bootstrapping the test data with 10 replicates.

Importantly, we found that the contribution of higher-order interactions measured in terms of percent epistatic variance increases substantially with distance to the data (Figure 5a, bottom panel). For nearby genotypes, the pairwise and the 4th-order model explain similar proportions of variance. However, as we move to more distant genotypes, the proportion of variance explained by higher-order interactions increases dramatically, such that at distance *>* 10, interactions of order 3-8 virtually account for all variance due to epistasis, translating to an additional 10 − 12% of total variance explained for test genotypes within each distance class.

We repeated the analysis using mean squared error (MSE) instead of *R*^2^ (Supplemental Figure 7) to account for the sensitivity of *R*^2^ to phenotypic variance within each distance class. Consistent with Figure 5a, we find that reduction of MSE of the additive model are driven primarily by pairwise interactions for genotypes near the training data, whereas improvements for distant genotypes are dominated by higher-order epistasis. We found similar data and phenotype distribution in the cgreGFP dataset (Figure 5b, first three panels).

The model predictive accuracy drops more rapidly than in the AAV2-Capsid dataset, such that the test *R*^2^ of the additive model falls below 0.35 for genotypes at mean distance *>* 9. The performance of the 8th-order epistatic transformer model also decreases with distance, but at a slower rate than the additive model. As a result, 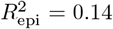 at distance 9, compared with 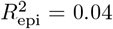 at distance ≤ 5. Similar to the AAV2 dataset, higher-order interactions become increasingly more important for distant genotypes, such that at distance ≥8, almost all the epistatic variance is due to interactions among more than 4 sites. Similar results were found when the analysis was repeated using MSE (Supplemental Figure 7).

In addition to the AAV2 and the cgreGFP dataset, we also apply the same analyses to four additional datasets (His-S2, -S5, -S12, and ppluGFP) that contain sufficient numbers of distance classes (Supplemental Figure 8). We observed that higher-order epistasis plays a moderate to significant role at greater distances in three of the four datasets, with the exception of His-S12, where epistatic effects were minimal or absent across all distances.

Together, our analyses show that higher-order interactions can be important for generalizing models trained on locally sampled data to distant regions of the sequence–function relationship. Furthermore, assessing the importance of higher-order epistasis using test data drawn from the same distribution as the training genotypes (e.g., in Figure 3) can be misleading, as the signal of higher-order epistasis may be much more prominent for distant genotypes. However, this pattern can be easily obscured because distant genotypes typically constitute a small proportion of the available data.

### Higher-order epistasis in a multi-peak fitness landscape

Having demonstrated that higher-order epistasis can help generalizing locally sampled fitness data to distant genotypes, in this section we provide another analysis of the importance of higher-order epistasis using a multi-peak fitness landscape. The data was derived from deep mutational scanning experiments for four fulllength GFP orthologs, avGFP, amacGFP, cgreGFP, and ppluGFP [20] (note that cgreGFP and ppluGFP2 were studied individually in the previous section). A single dataset was generated by merging the four GFP datasets based on the multiple sequence alignment of the four WT sequences [20]. The four proteins exhibit moderate to very high sequence divergence ranging from 18% (between avGFP and amacGFP) to 83% (between ppluGFP2 and amacGFP) (Figure 6a). This strong divergence allows us to circumvent the limitations of locally sampled data in revealing higher-order epistasis, which can be masked by strong correlations in mutational and pairwise interactions due to similarities among genetic backgrounds.

**Figure 6.**
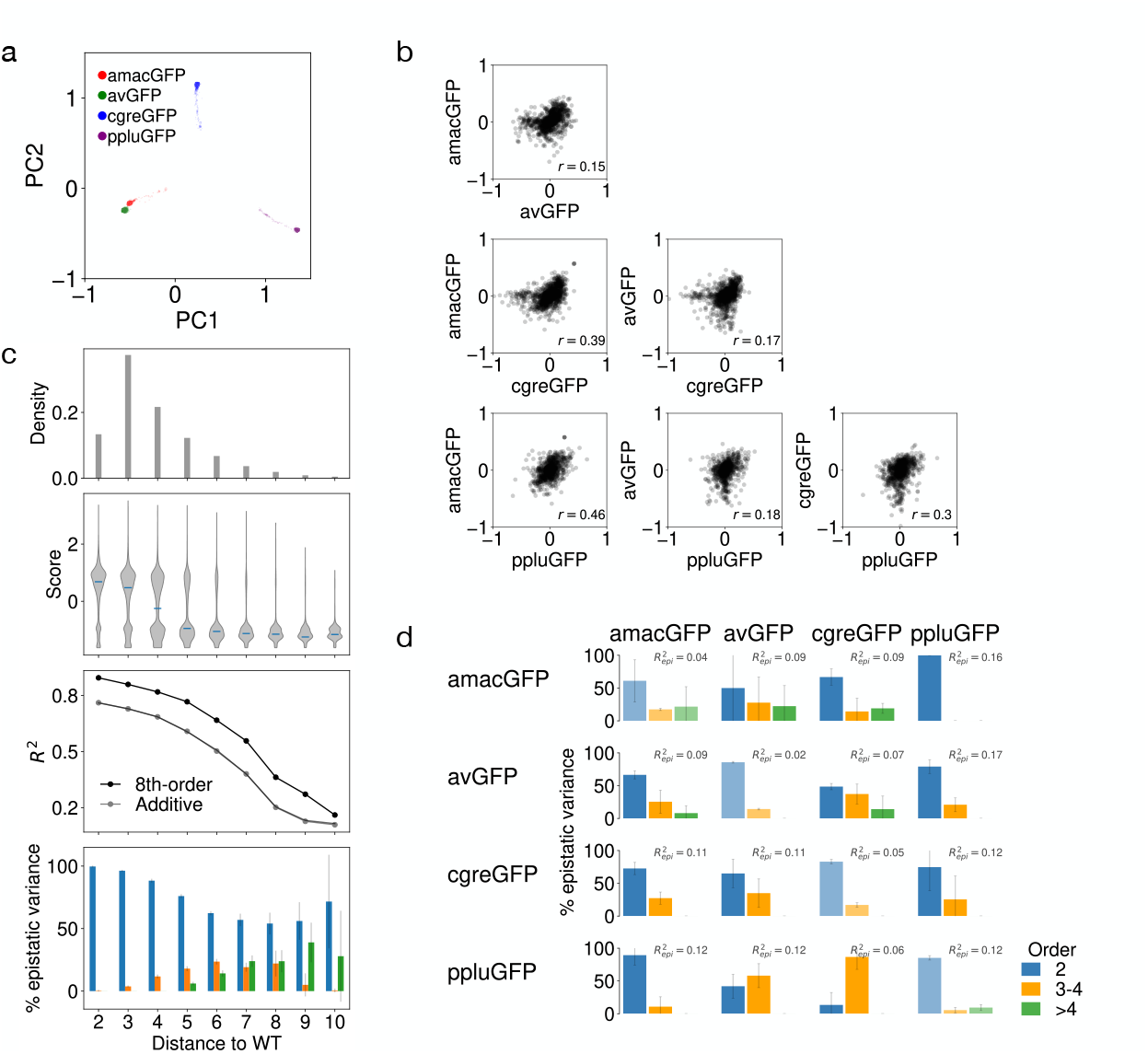
Higher-order epistasis in a multi-peak fitness landscape, consisted of four green fluorescent protein (GFP) orthologs (avGFP, amacGFP, ppluGFP2, cgreGFP). a. PCA coordinates for the one-hot embeddings of the all protein genotypes. Genotypes are highly concentrated around the four wild types (WT), which exhibit varying degrees of sequence divergence. b. Scatter plots of shared mutational effects among the four GFPs, fit using separate additive model with a shared sigmoid activation function. c. Higher-order epistasis allows better generalization to distant regions in sequence space. Models were fit use single and double mutant data for all GFPs. Models were tested in different distance classes, each containing all genotypes at given Hamming distance to their corresponding WT sequence. Error bars represent 1 standard deviation calculated by resampling random 90% of the test genotypes with 10 replicates. d. Higher-order epistasis allows better generalization across local peaks. Rows: GFP orthologs used to train the models. Columns: GFP orthologs used to test performance of the trained model. 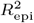 at top right corner of each panel measures the proportion of total variance explained by specific epistasis among up to 8 positions. Error bars correspond to 1 standard deviation across three model replicates with different training and test genotypes.

Prior analyses of this dataset [20] identified broad similarities among the four fitness peaks, including strong threshold effects of mutational distance on fluorescence and distinct effects of mutations of surface versus buried residues, as well as substantial differences in peak sharpness and the role of local epistasis. To provide additional coarse characterization of the (dis)similarity among the four local peaks, we directly compare the effects of shared mutations for all pairs of GFPs in Figure 6b . Here, the mutational effects were derived from individual additive models with a shared sigmoid activation function to ensure the inferred mutational effects are on the same scale. We found that mutational effects overall have low to moderate correlation between GFP orthologs, with Pearson *r* ranging between 0.15 to 0.46. Interestingly, sequence divergence between orthologs has no effect on similarities in mutational effects. For example, the strongest correlation (*r* = 0.46) is between amacGFP and ppluGFP2, which exhibit very high sequence divergence (83%).

With this basic understanding of the divergence between the GFP orthologs, we proceed to quantify the importance of higher-order epistasis in this multi-peak landscape. We first performed a baseline task by fitting the pairwise, 4th-order, and 8th-order epistatic transformer models to 80% of randomly sampled variants from all four GFP orthologs. We see that fitting epistasis up to 8th order leads to an overall 12% improvement in test *R*^2^ (Supplemental Figure 9). In particular, we found that pairwise interactions are the predominant form of epistasis, while higher-order epistasis accounts for 15% of the total epistatic variance in the test genotypes. This result suggests that pairwise interactions are capable of capturing most of the phenotypic variances among distant local fitness peaks, potentially by accounting for the divergence in local mutational effects (e.g. Figure 6b).

Next, we performed a more stringent task by training the model on all single and double mutants across the four orthologs and assessing performance of the pairwise and higher-order epistatic transformer models on distant, higher-order mutants. Figure 6c shows that prediction accuracy for both the additive model and epistatic transformer with 3 epistatic MHA layers declines with distance from the fitness peaks. While the total epistatic variance 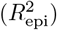 remains relatively constant, higher-order interactions contribute an increasing fraction of this variance for more distant genotypes, accounting for roughly half of the epistatic variance for sequences at Hamming distance 7-9 to their respective orthologs. This result suggests that, although the training data contain only local mutational effects and pairwise interactions, the epistatic transformer can approximate fine-scale higher-order epistasis within individual peaks using coarse-grained higher-order interactions encoded in the differences in local effects across highly divergent genetic backgrounds.

For the final task, we fit epistatic transformer to random training genotypes for each GFP and examine how models fit to individual local peaks can extrapolate to other orthologous GFP landscapes. Our hypothesis is that the higher-order epistasis learned by our model when fit to one local fitness peak will help predict how locally observed mutational effects and pairwise epistasis generalize to highly divergent, novel backgrounds, thus enabling better prediction. For each GFP ortholog, we trained the models using 50% of randomly chosen genotypes and a validation dataset consisting of the most distant 30% of the remaining genotypes. We then assessed model generalizability by predicting fitness for random test genotypes sampled from each of the four GFP orthologs.

We first note that the higher-order models explain low to moderate proportions of variance for test genotypes within the same local peak (Figure 6d, diagonal panels). In contrast, when predicting genotypes from different orthologous fitness peaks (Figure 6d, off-diagonal panels), higher-order interactions frequently make substantial contributions to the total epistatic variance, sometimes matching or exceeding the percent epistatic variance explained by the pairwise model (e.g., a model trained on ppluGFP and evaluated on avGFP and cgreGFP).

Together, our results show that higher-order epsitasis is important in determining the structure of this multi-peak GFP landscape. Furthermore, higher-order epistasis can be learned from both the local and the global scale to enable better out-of-distribution predictions.

### Architecture of specific epistasis for GRB2-SH3 abundance

Although the epistatic transformer architecture allows us to probe the relative importance of higher-order versus pairwise epistatic interactions, it does not directly review the actual biological interactions captured by the epistatic MHA mechanism. In this section, we provide an exploratory analyses using two principled methods in order to gain additional insights into the genetic architecture of epistasis. We use GRB-abundance dataset as an example, as it shows the largest contribution of higher-order epistasis among the chosen datasets. Furthermore, we perform all analyses on the 8th-order epistatic transformer model fit to 80% of training data, and apply our methods to the latent phenotype *ϕ* (before the sigmoid activation function) to remove the influence of the global nonlinearity.

We first employed a method to calculate site-specific marginal epistatic effect (see Supplement for mathematical details). In particular, this method allows us to estimate, for a focal site *l*, how much of the total variance due to *k*-order epistasis is explained by all *k*-way interactions involving *l*. It can be shown that this quantity is proportional to the sum of interaction coefficients involving *l* under certain regularity conditions, such that for *k* = 3 the marginal epistatic effect for *l* is

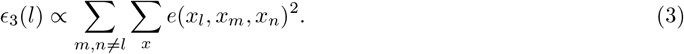

Therefore, this quantity is useful for identifying positions enriched for pairwise or higher-order epistasis. An advantage of this method, compared with standard explainable AI approaches such as DeepLift [63] and DeepSHAP [64], is that it does not require arbitrarily chosen reference sequences, whilst allowing us to study the contribution of individual sites to specific orders of epistasis.

In Figure 5a, we summarize the marginal additive, pairwise, three-way, and four-way epistatic effects across the 34 mutagenized sites for GRB2-SH3. We first see that the sites roughly exhibit a two fold variation in marginal epistatic effects across orders. Furthermore, we observed that the site-specific marginal effects are consistent for pairwise, three-way, four-way interactions (Pearson *r >* .9), such that a site important for pairwise epistasis is likely to enrich for three- and four-way interactions. Interestingly, the marginal epistatic effects appear to have low correlation with the additive contributions (Pearson *r <* 0.2). For instance, positions 4 and 8 both have strong additive effects and explain around 10% of the total additive variance, but are among the least epistatic sites.

Overlaying the pattern of marginal epistatic effects on the 3D structure (Figure 7b), we found that positions enriched for epistatic interactions (e.g. site 20-46) are spread across three beta sheets and part of a loop region connecting this segment to the N-terminal beta sheet. In contrast, sites on the N-terminal and C-terminal *β* sheets exhibit much lower levels of epistasis. This result is not surprising as the central *β*-sheet cluster forms a tightly packed network of hydrogen bonds which likely constrains local movements and promotes higher-order coupling, whereas the more solvent-exposed and flexible terminal regions can accommodate substitutions with minimal disruption to the fold, leading to weaker epistasis.

**Figure 7.**
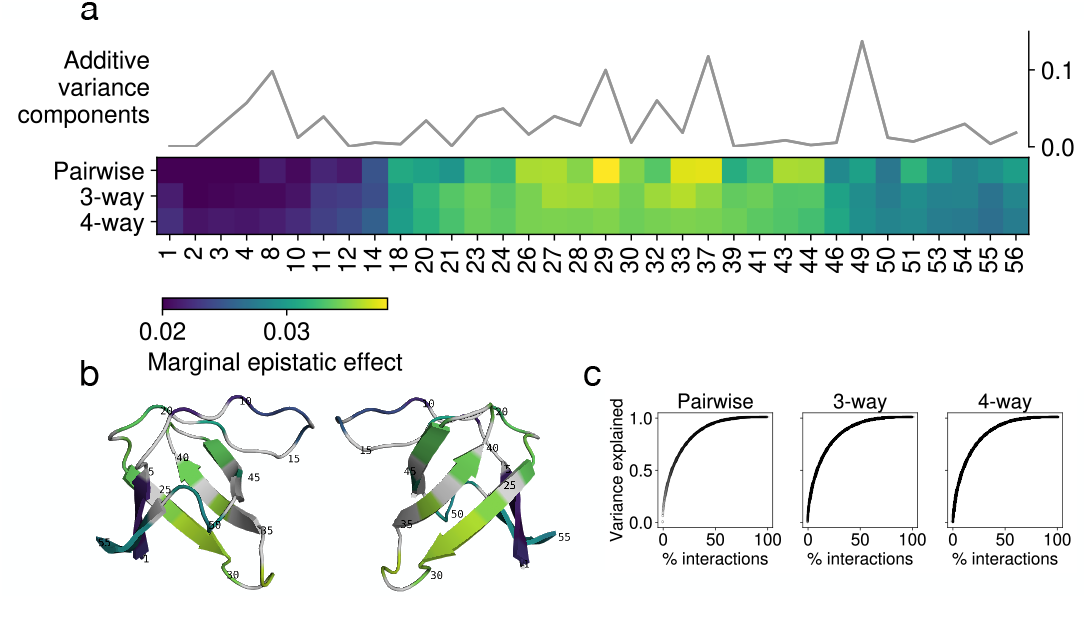
Genetic architecture for epistasis of GRB2-SH3 abundance, inferred using a 3-layer epistatic transformer model fit to the ‘GRB-abundance’ dataset. a. Position-specific additive and marginal epistatic effects. Additive effects were calculated as the fraction of total additive variance explained by each position. Marginal epistatic effects of order *m* (*m* = 2, 3, 4) for position *i* were calculated as the fraction of total epistatic variance of order *m* explained by all *m*-way interactions involving site *i*. Effects were normalized to sum to 1 for row to assist in visualization of the results. Results were derived by averaging over 5 replicates over random subsamples of all available data. b. Marginal three-way epistatic effects mapped to the 3D-structure of GRB2-SH3 (pdb: 1GCQA). Gray corresponds to unmutagenized positions in the experiment. c. Sparsity of epistasis. Each panel shows the fraction of total pairwise, three-way, or four-way variance explained by the top percentage of interacting sites. For example, the leftmost panel shows the top 50% all pairs of sites explain nearly all of the variance due to pairwise epistasis.

We next introduce a second statistic for quantifying the marginal interaction strength among particular positions. For instance, given sites *l, m, n*, our method quantifies the proportion of total phenotypic variance due to pure three-way interactions among these three positions, by averaging over all genetic backgrounds, whilst removing the influence of both lower-order and higher-order epistasis (see Supplement).

With this, we quantify the marginal epistatic effects for all subsets of 2, 3, and 4 sites. In particular, we ask if specific epistasis exhibit a sparse pattern across interaction orders, i.e., if a small number of interactions account for most of the epistatic variance. In Figure 7c, we show the proportion of total variance due to pairwise, three-way, and four-way epistasis explained by the top percentage of interacting sites. We can see that all three curves saturate relatively fast, such that the top 10% of the interacting subsets explain 50% of the variance due to pairwise, three-way, or four-way interactions, revealing moderate sparsity in the epistatic structure in the GRB2-SH3 abundance fitness landscape.

## Discussion

We developed a novel neural network-based method capable of detecting higher-order epistasis in full-length protein sequence-function relationships. The study of epistasis in high-throughput protein datasets has been historically hindered by the combinatorial complexity of fitting higher-order interaction terms, which makes existing methods applicable only to protein sequence-function datasets with at most 3-4 positions mutagenized to every other amino acid in a combinatorially complete manner. By contrast, our epistatic transformer model fits epistatic interactions implicitly through a neural network, thus can be easily applied to study epistasis among hundreds of amino acid positions.

We applied our method to 10 protein sequence-function datasets. Our results showed that higher-order epistasis can have subtle to substantial effects in determining the overall structure of protein fitness landscapes. The strongest effect of higher-order epistasis was observed in the abundance data for the SH3 domain of the protein GRB2, where epistasis among ≥ 3 sites explains 15% of the total variance in test genotypes, representing nearly two thirds of the total variance due to epistasis. This finding contrasts with a previous study [31], which discovered no substantial contribution by higher-order epistasis in 20 protein sequence-function relationship datasets. This discrepancy may be explained by the different scales of the study datasets. While Park et al. [31] exclusively focused on combinatorially complete sequence-function data for small protein sequence space, we have predominantly used locally sampled sequence-function data embedded in much larger genotypic spaces, where higher-order epistasis likely plays more important roles for combinatorial and biophysical reasons [65]. Our finding suggests that while some protein sequence-function relationship may be simple and explainable by models only considering additive effects, pairwise epistasis, and non-specific epistasis, higher-order specific epistasis can play a critical role in other cases and thus must be properly considered to fully understand the structure of protein sequence-function relationships.

It is also noteworthy that, in most of the other datasets we analyzed, the contribution of higher-order interactions is relatively modest when measured in terms of the total variance they explained, alignining with the findings of Park et al. [31]. One possible explanation for this seemingly limited contribution of higher-order interactions is that these protein-sequence function relationships are truly devoid of higher-order interactions. Alternatively, higher-order epistasis may be present but obscured by experimental limitations, potentially due to the choice of mutagenized sites that do not interact strongly, or the limited scope of local sampling which does not ensure adequate statistical power to detect high-order interactions.

Importantly, even when datasets show little evidence of higher-order epistasis in summary statistics like *R*^2^, such interactions can still play crucial roles in determining the phenotypes of a minority of genotypes with idiosyncratic behavior [66]. We demonstrated this by evaluating the importance of higher-order epistasis for making predictions for out-of-distribution genotypes. In particular, we showed that higher-order effects can be essential for predicting phenotypes in sparsely sampled regions of the sequence space, and for generalizing from data concentrated in one local GFP fitness peak to other, more distant orthologs. These findings suggest that coarse-grained metrics such as *R*^2^ may indeed fail to fully capture the functional significance of higher-order epistasis. Relying on such summary statistics to guide protein engineering or interpret evolutionary patterns can therefore be misleading, particularly when extrapolating beyond the range of observed data into unexplored regions of the sequence space.

Additionally, it may also be tempting to ask if the variation in the prevalence of epistasis can be explained by different properties of the proteins we studied, such as protein families, enzymatic functions, and structural features. At present, we cannot address this question, because the number of high-order, combinatorial DMS datasets are still very limited. As these types of experiments become more common, revisiting this question will surely be warranted.

Our results may present a challenge to researchers in protein engineering and evolution as they suggest that higher-order epistasis cannot be altogether ignored when studying protein-sequence function relationships. We want to point out that the extent to which researchers should ‘worry’ [28] about higher-order epistasis depends on the particular application. For example, based on the evidence of the Park et al. study [31], higher-order epistasis is likely to be less prevalent when only a small number of sites are mutagenized. Similarly, higher-order epistasis may also play less important roles in determining the structure of local sequence-function relationships. However, if the researcher is interested in how large numbers of mutations across the whole protein combine to determine function, then higher-order epistasis should be properly incorporated. Reassuringly, we found that higher-order epistasis can be effectively modeled based on moderate-size data to enable a faithful construction of the sequence-function relationship.

## Methods

### Epistatic transformer architecture

In this section, we provide a detailed description of the architecture of the epistatic transformer model in Figure 1. Our input *x* consists of tokenized protein sequences, i.e. *x* = *x*_1_, *x*_2_, …, *x*_*L*_, where *x*_*l*_ is an integer between 0 and *a*−1 (*a* is the alphabet size, which depends on the study dataset). We used a special positional encoding to transform the raw input sequence such that for the new sequence 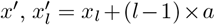. That is, we essentially treat amino acids from different positions completely differently. This is different from typical protein language modeling where a continuous positional encoding is added to the amino acid embeddings [67– 69], such that the identities of the amino acids are still largely preserved to allow the model to learn general rules of interaction among residues while informed by residue positions when trained on large number of natural protein sequences. In contrast, here we aim to have the maximum flexibility to allow the same amino acid combinations to have different interactions at different positions, in order to learn a model that well approximates the parameterization in Eq. 2.

The tokenized sequence *x*^*′*^ is passed through an embedding layer to generate position-wise continuous representations 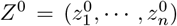, where each position consists of *d* hidden dimensions. The embeddings are then passed through *M* layers of modified multi-head attention (epistatic MHA) to model epistatic interactions among sites. Our epistatic MHA layer is built on the standard MHA layer, but with some key differences. Specifically, the input *Z*_*m*_ ∈ ℝ^*d×L*^ from MHA layer *m* is first used to generate the query and key tensors for *H* attention heads in layer *m* + 1, such that for head *h*, the query and key tensors are

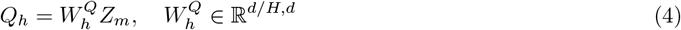

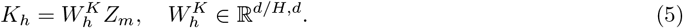

Importantly, the value tensor *V* is directly generated from the raw embedding *Z*_0_, instead of *Z*_*m*_

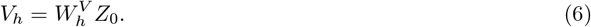

In Supplemental Methods, we show how this modification is essential to ensure that *M* layers of epistatic MHA gives rise to models that strictly contain specific epistasis of order up to 2^*M*^ . Next, for head *h*, we perform a modified scaled dot attention (Figure 1)

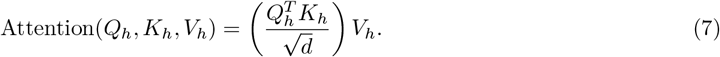

The key difference between this equation and the standard scaled dot attention is the lack of a softmax normalization, which is used to convert the raw attention scores obtained from the dot product of the query and key vectors into a probability distribution over the input positions. Since in the standard scaled dot attention the softmax operation would be applied across positions for each embedding dimension, it acts similarly to a global epistasis nonlinearity. This would cause the hidden nodes in the model to contain interactions of all orders, which is unsuitable for our purpose of only fitting specific epistasis up to a particular order. Using our modified scaled dot attention, the output from each head is then concatenated and passed through a linear layer to generate the output *Z*_*m*+1_

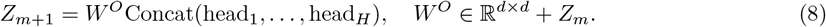

The output of the final *M* th MHA layer is then flattened and converted to a scalar *ϕ*

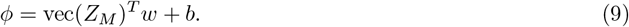

The scalar *ϕ* can be interpreted as a hidden phenotype that only contains additive effects and effects due to specific interactions. Note that our construction of the model architecture makes sure that *ϕ* only contains specific interactions of order up to 2^*M*^ . The scalar *ϕ* is then passed through a sigmoid function mapping the hidden phenotype to the measurement scale to account for any non-specific epistasis

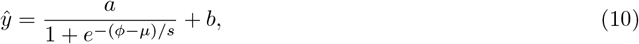

where *a, µ, s*, and *b* are trainable model parameters, which allow us to shift and scale the standard sigmoid function. We also experimented with more complex nonlinear activation functions parameterized by a sum of independent sigmoid functions. We did not observe improvement with this method over Eq. 10 (Supplemental Figure 1). Thus, we trained all models using Eq. 10 as the global epistasis function. Another key difference between our model and standard transformer models is the lack of LayerNorm and position-wise feed-forward networks, as both will introduce undesired global epistasis effect in the hidden layers.

### Model training

For all datasets studied in the main text, we train the epistatic transformer model with 1, 2, and 3 layers of epistatic MHA, corresponding to models with specific epistasis among up to 2, 4 and 8 sites. In addition to the epistatic models, we also trained an additive + global epistasis model to benchmark the importance of specific epistasis. The additive model uses the same sigmoid activation function in Eq. 10 to model nonspecific epistasis. However, its hidden phenotype *ϕ* was calculated as a simple weighted sum of the flattened one-hot embeddings of the input sequence, which was used to model the sum of additive effects of all mutations.

As mentioned in the main text, we use Optuna [60] to optimize hyperparameters including hidden dimension (*d*), number of train epochs, batch size, and dropout rate. Optuna uses a Bayesian optimization algorithm called Tree-structured Parzen Estimator to model the history record of trials to determine which hyperparameter values to use for the next iteration. We use the *R*^2^ value on a validation dataset to guide 200 Optuna optimization steps for all models. In our preliminary study, we found that learning rate and the number of attention heads have little effect on the final model performance, we therefore used 4 attention heads and the Adam optimizer with fixed learning rate (0.001) to train all models. All code was written in PyTorch v2.2.0. Individual models were trained on one NVIDIA Ampere A100 GPU with 80GB memory.

### Selection of experimental datasets

We used the following criteria for selecting datasets for detecting higher-order epistasis by fitting our epistatic transformer model. First, the dataset must contain high-throughput measurements for proteins, and ideally embedded in a large sequence space to allow potentially non-significant contribution by higher-order epistasis. Second, the dataset must contain substantial numbers of higher-order mutants, i.e. genotypes with at least three mutations relative to the reference (WT) sequence, to ensure enough statistical power for detecting higher-order epistasis. We found that combinatorial mutagenesis datasets satisfying these criteria are rare compared with the large body of studies that measure single mutant and/or double mutant effects. Our research resulted in 7 publications. However, initial fitting of our models to datasets from two publications [70, 71] resulted in very low model performance (*R*^2^ *<* .1 for all models), which were subsequently discarded. The final datasets are listed in Table 1, containing data from five publications.

Three publications contain measurements for multiple proteins or functions. We used all three experiments conducted for the SH3 domain of the growth factor receptor-binding protein GRB2 by Faure et al. [23]. The His3 publication [22] contains fitness measurements for combinatorial variants in 12 non-overlapping segments tiling the His3 protein. We chose segment 2, 5, and 12 for our study as these three segments showed the lowest *R*^2^ when we fit an additive + sigmoid model, thus allowing for more room of improvement by the epistatic models. The GFP publication [20] contains fluorescence data for three GFPs (amacGFP, ppluGFP, and cgreGFP), complemented by a fourth GFP (avGFP) [51] from an earlier study. For examining our model performance on random training data, we only used the cgreGFP and the ppluGFP datasets. These four datasets were combined using the multiple sequence alignment of the four wild type sequences to generate the a multi-peak sequence-function relationship dataset.

## Data Availability

Computer code to replicate all analyses can be found at https://github.com/juannanzhou/EpistaticTransformer.

## Acknowledgement

We thank David McCandlish for thoughtful comments and suggestions. Research reported in this publication was supported by the National Institute of General Medical Sciences of the National Institutes of Health under award number R35GM154908, and University of Florida College of Liberal Arts & Sciences.

## Author contributions

J.Z planned research; P.S and J.Z performed research; and J.Z wrote the paper.

