## Supplement for "An interpretable neural network unveils higher-order epistasis in large protein sequence-function relationships"

#### Contents

**1 Supplemental Figures** **2**

**2 Supplemental Methods** **12**

2.1 Analyses of simulated data . . . . . 12

2.2 Proof highest-order of epistasis in epistatic transformer scales as $2^M$ with attention layers $M$ 12

2.3 Calculation of marginal epistasis . . . . . 14

### 1 Supplemental Figures

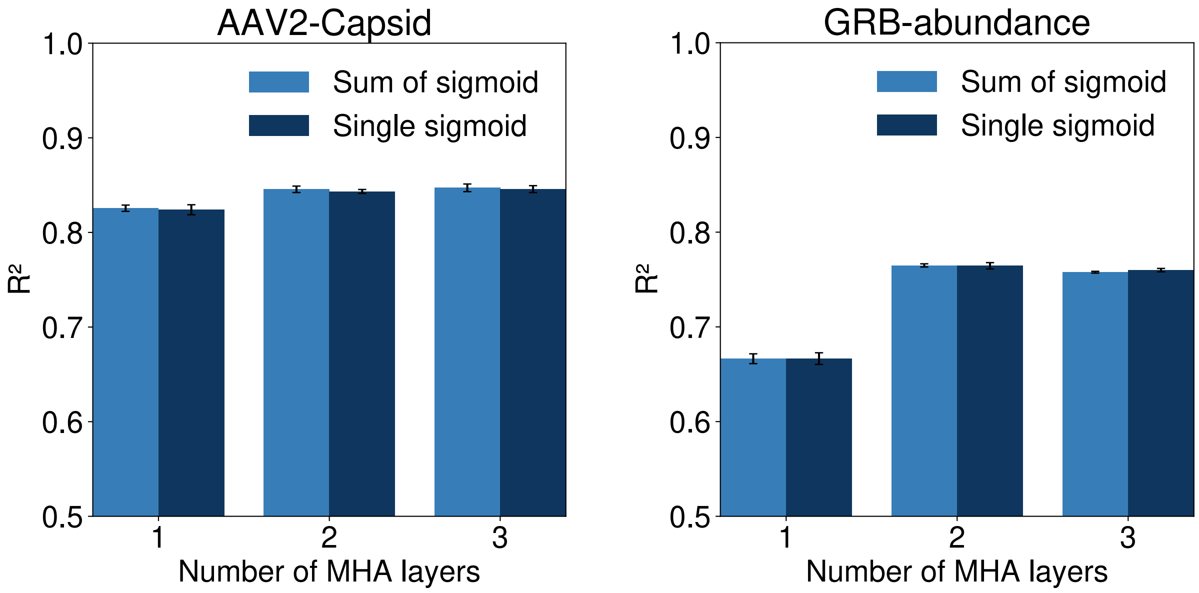


Supplemental Figure 1: Single sigmoid activation function is sufficient for modeling global epistasis. Epistatic transformer model with $M = 1, 2, 3$, layers of attention were trained on 50% of random genotypes for the AAV2 and GRB2-abundance datasets. The final activation function take the form of a single sigmoid activation function with the form $\sigma(z) = \frac{a}{1+e^{-(z-\mu)/s}} + b$ vs. sum of 3 separate functions of this form.

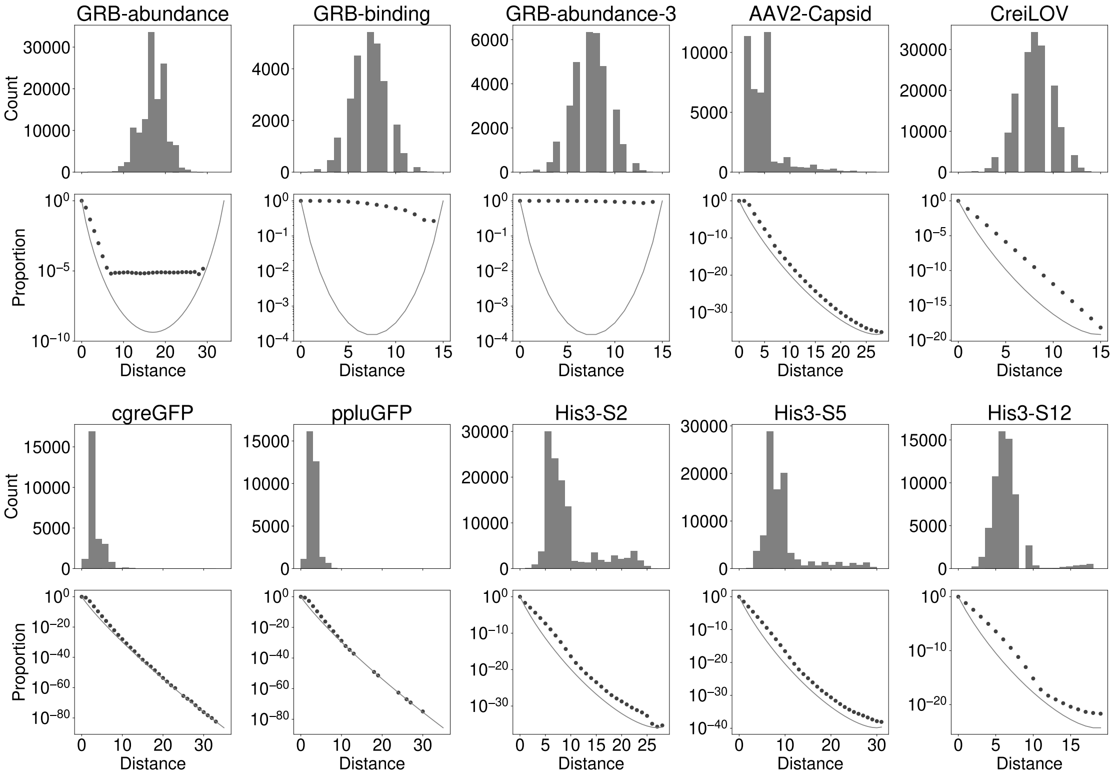


Supplemental Figure 2: Data distribution for the protein mutagenesis datasets analyzed in this paper. For each dataset, the top panel is the histogram for variant counts at different Hamming distances to the reference (wild type) sequence. The bottom panel shows the proportion of measured variants for the distance classes (dotted lines). Solid lines show the inverse of the size of the distance class, corresponding to the proportion when only one variant is measured in the class.

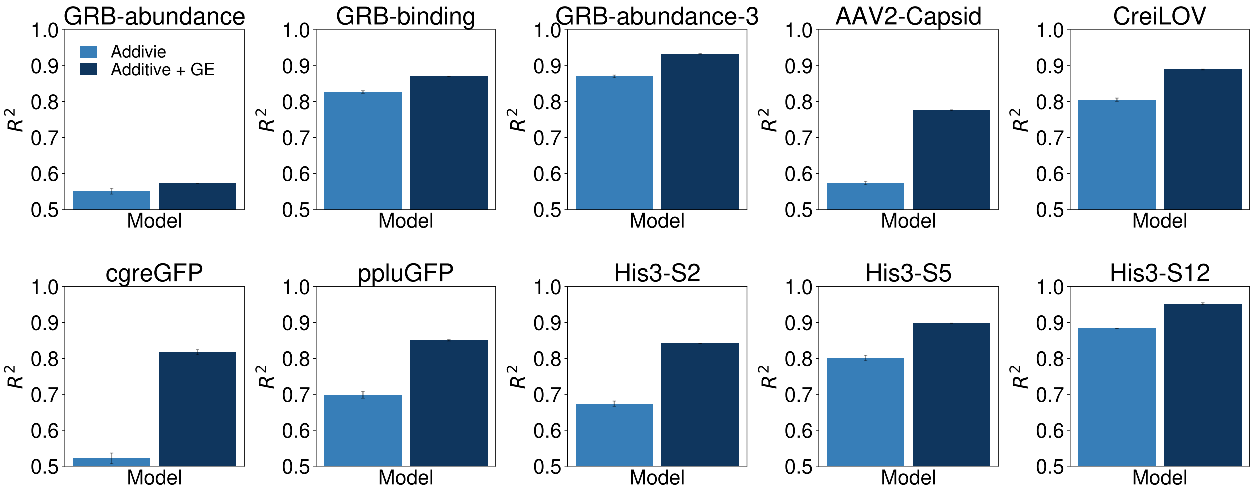


Supplemental Figure 3: Importance of incorporating global epistasis to the additive model. Additive models without and with the sigmoid activation function were fitted to 80% of random training data in all 10 datasets. $R^2$ scores were calculated on random test genotypes. Error bars represent 1 standard deviation calculated based on 3 replicates.

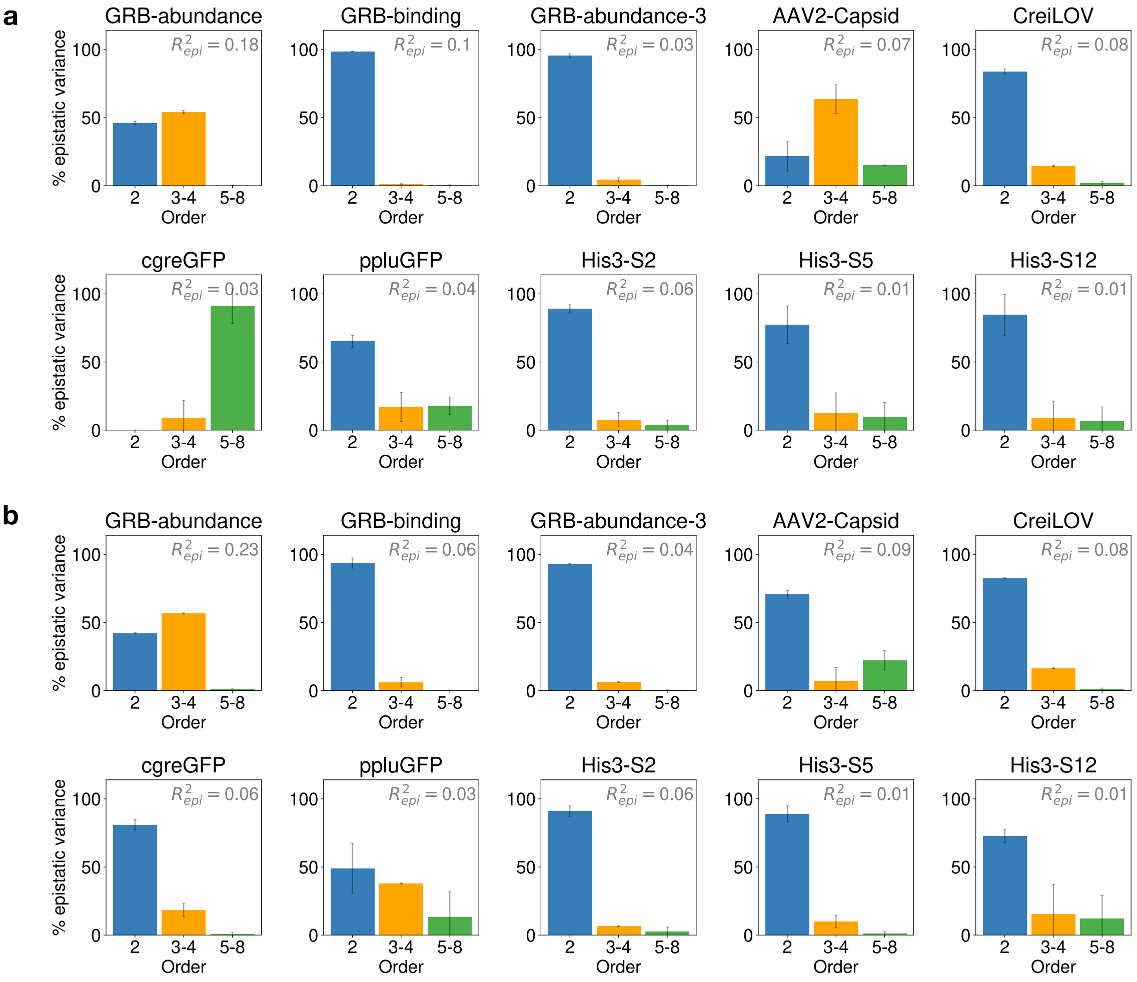


Supplemental Figure 4: Importance of pairwise and higher-order specific epistasis in 10 experimental protein-sequence function datasets. For each dataset, pairwise, 4th-order and 8th-order models were fitted using epistatic transformer with 1, 2, and 3 layers of epistatic attention. All models contain a final nonlinear activation function mapping a scalar value to the measurement scale for modeling non-specific epistasis. Models were fitted to 20% (a) and 50% (b) of training data generated by randomly sampling all available data. In each panel, the number $R^2_{\text{epi}}$ on the upper right corner is equal to the proportion of total variance in the test data explained by all orders of specific epistasis, equal to the difference between the $R^2$ of the 8-th order epistatic transformer model and the additive model. Importance of epistatic interactions of different orders is measured by percent epistatic variance, equal to the gain in $R^2$ by fitting an additional layer of MHA, normalized by $R^2_{\text{epi}}$. Error bars represent 1 standard deviation calculated from 3 replicates.

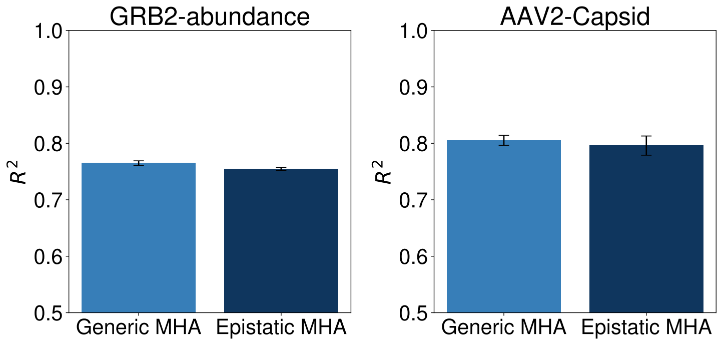


Supplemental Figure 5: Performance comparison between standard transformer with generic multi-head attention (MHA) and epistatic transformer with epistatic MHA. Models were fit to 20% of randomly sampled genotypes for the AAV2 and GRB2-abundance dataset. Both models have three layers of attention. Error bars represent 1 standard deviation, calculated from 3 replicates.

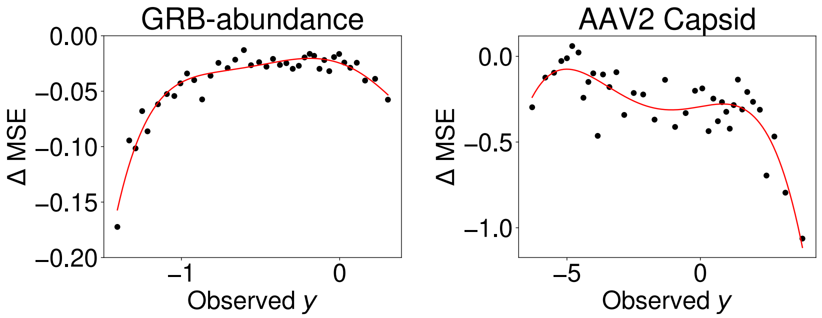


Supplemental Figure 6: Improvement of model performance by the high-order epistatic model is achieved throughout the whole range of observed phenotypes for both the GRB-1 and AAV2 Capsid datasets. y-axis: Differences in mean squared error (MSE) on test genotypes between the 8th-order and pairwise epistatic transformer models. Negative values indicate better performance by the 8-th order model. ΔMSE was calculated for different bins of test genotypes sorted by their phenotypic ($y$) values. Red curve is a polynomial function fit to the data points.

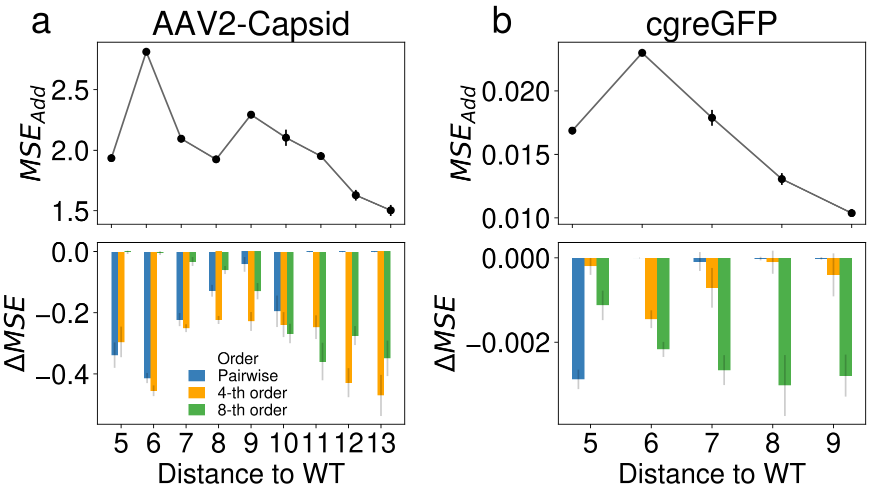


Supplemental Figure 7: Importance of higher-order epistasis in predicting phenotypes for distant genotypes in the AAV2-Capsid (a) and the cgreGFP (b) datasets, measured in terms of test mean squared error (MSE). Genotypes are binned to discrete distance classes by their mean Hamming distances to the training data, which consist of 20% of randomly sampled genotypes. For both datasets, we retain only distance classes where the additive model has a test $R^2 > 0.3$. Top panel: Test MSE under the additive model for genotypes at different distance classes. Bottom panel: improvement in prediction performance by epistatic transformer models of different orders. $\Delta$ MSE is calculated as the difference in test MSE between the current model and the precedeing, less complex model. All metrics were calculated for models fitted to one training sample consisting of 20% of randomly sampled genotypes. Error bars represent 1 standard deviation calculated by resampling the test data with 10 replicates.

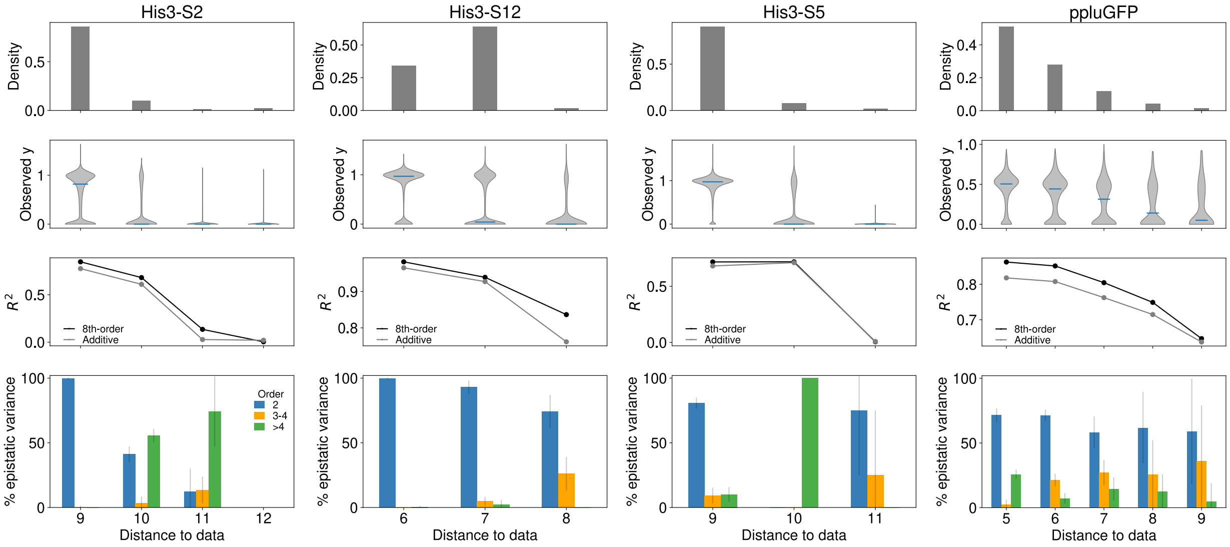


Supplemental Figure 8: Importance of higher-order epistasis in predicting phenotypes for distant genotypes in 4 datasets. For each dataset, genotypes are binned to discrete distance classes by their mean Hamming distances to the training data, which consist of 20% of randomly sampled genotypes. Top panel: Distribution of mean Hamming distance in the randomly sampled test data. Second panel: Distribution of phenotypic scores for each distance class. Third panel: Test $R^2$ under the additive model and the 8-th order epistatic transformer for genotypes at different distance classes. The gap between the two curves is equal to $R^2_{\mathrm{epi}}$ for each distance class. Bottom panel: importance of specific pairwise and higher-order epistasis at different distance classes, measured by percent epistatic variance, equal to the gain in $R^2$ by fitting an additional layer of MHA, normalized by $R^2_{\mathrm{epi}}$. Error bars represent 1 standard deviation calculated by bootstrapping random 90% of the test genotypes.

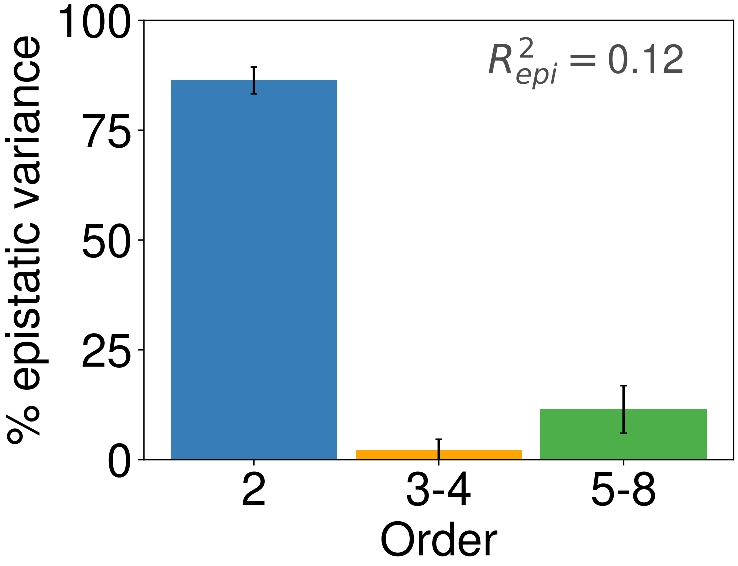


Supplemental Figure 9: Predictive performance of epistatic transformer models with increasing complexity to the aligned GFP dataset consisted of fluorescent levels for variants belonging to four GFP orthologs (amacGFP, avGFP, cgreGFP, and ppluGFP). Models were trained on 80% of randomly sampled genotypes. Y-axis corresponds to the percentage of epistatic variance in the test data explained by different orders of interactions. Error bars correspond to 1 standard deviation calculated from 10 randomly samples of test genotypes.

### 2 Supplemental Methods

#### 2.1 Analyses of simulated data

In this section, we describe how we generated the data used in Figure 2 of the main text, which is used for characterizing the behavior of the epistatic transformer model. We simulated data for a sequence space with 13 sites, each with two alleles. The sequence-function relationship was generated using this equation

$$\phi(x) = \sum_{i=1}^{13} \beta_i x_i + \sum_{1 \le i < j \le 13} \beta_{ij} x_i x_j + \sum_{1 \le i < j < l \le 13} \beta_{ijk} x_i x_j x_l + \cdots , \tag{1}$$

where the dots symbolize higher-order interactions among up to all 13 positions. Specifically, $x_i = 1$ or $-1$, such that $x_i x_j$, $x_i x_j x_l$ and their higher-order analogs form an orthogonal bases for the space of all sequence-function relationships. Consequently, the $\beta$ parameters correspond to the Walsh-Fourier coefficients.

We sampled the $k$th-order Walsh coefficients independently from a zero mean Gaussian distribution where the variance (namely $\lambda_k$) was parameterized by an exponential decay function, such that $\lambda_k = e^{-1.1\times k}$.

To calculate the variance components ($V_k$, equal to the proportion of total variance explained by $k$-order interactions) of the simulated data, we use the formula

$$V_k = \frac{a^{-L} \boldsymbol{\phi}^T P_k \boldsymbol{\phi}}{\mathrm{Var}(\boldsymbol{\phi})}, \tag{2}$$

where $P_k$ is the projection matrix to the subspace spanned by all pure $k$th-order interactions, and is defined element-wise by the Krawtchouk Polynomial

$$P_k(x, x') = W_k(x, x') = W_k(d) = a^{-L} \sum_{q=0}^{k} \binom{d}{q} \binom{L-d}{k-q} (-1)^q (a-1)^{k-q}, \tag{3}$$

where $d$ is the Hamming distance between the two sequences $x$ and $x'$. Therefore, $\boldsymbol{\phi}^T P_k \boldsymbol{\phi}$ is equal to the squared norm of the $k$th-order component of $\phi$. Consequently, the numerator in Eq. 3 is the variance of the projected vector, making Eq. 33 the $k$th-order variance component of $\phi$. In Figure 2, we also used a related quantity, the cumulative variance components ($V_k'$) to examine if out-of-sample $R^2$ of the trained model can be used to infer the amount of epistasis in the data explained by interactions up to the 2, 4, and 8-order interactions. $V_k'$ is equal to the sum of variance explained by all orders of interaction not higher than $k$. Thus, we can simply calculate it as

$$V_k' = \sum_{i=1}^{k} V_k. \tag{4}$$

All models were trained using Optuna hyperparameters resulted from 100 iterations on 90% of randomly sampled data. The (cumulative) variance components of the final trained model as well as the intermediate model shown in Figure 2b and 2c were calculated using Eqs. 2 and 4.

#### 2.2 Proof highest-order of epistasis in epistatic transformer scales as $2^M$ with attention layers $M$

In this section, we provide a proof for the key property of the epistatic transformer model that it can fit specific epistasis up to order $2^M$ in the latent phentoype $\phi$, with $M$ epistatic multihead attention layers.

We first begin by proving the following lemma.

**Lemma 2.1.** *The output of the mth multihead attention layer for position l, $z_l^m$ can be expressed as*

$$z_l^m = \sum_{\mathbf{i}} f(x_l, x_{i_1}, x_{i_2}, \cdots), \tag{5}$$

*where the sum is over all multi-indices* $\mathbf{i}$ *such that* $|\mathbf{i}| \le 2^m - 1$, *each containing a subset of all positions between* 1 *and* $L$. $f$ *is some function that depends on* $l$, *positions* $\mathbf{i}$, *and the model parameters* $\theta$.

*Proof.* We prove this by induction. Recall that $z_l^0$ is the input embedding generated using the the token $x_l$. It is easy to see that when $m = 0$, the lemma holds trivially. Next, assume that Eq. 5 is true for $m \geq 0$. We need to show that Eq. 5 is also true for $m + 1$. To begin, recall that the output of the $m + 1$ MHA layer by one of the attention head $h$ before the linear transformation is

$$u_l^{m+1,h} = \sum_{l'} \left( \frac{(W_h^Q z_l^m)^T W_h^K z_{l'}^m}{\sqrt{d}} \right) z_{l'}^0, \tag{6}$$

where $z_l^m$ is the output of the $m$th MHA layer for position $l$ and $z_l^0$ is the raw amino acid embedding that only depends on $x_l$ and no other positions. Note that we do not apply a softmax function across all positions $l'$ to generate normalized attention weights as regular transformers.

This simplification allow us to express Eq. 6 as

$$u_l^{m+1,h} = \sum_{l'} \langle z_l^m, z_{l'}^m \rangle \, z_{l'}^0, \tag{7}$$

Here $\langle z_l^m, z_{l'}^m \rangle$ is some function that is bilinear in $z_l^m$ and $z_{l'}^m$. It is easy to see this is the case as the mapping from the input vectors $z_l^m, z_{l'}^m$ to the query or key vector is linear, and that the calculation of the attention weight is a dot product (due to the lack of the softmax operation). We can then bring Eq. 5 into Eq. 7. Using the bilinear property of the $\langle , \rangle$ function, Eq. 7 equation becomes

$$\begin{aligned} u_l^{m+1,h} &= \sum_{l'} \langle \sum_{\mathbf{i}} f(x_l, x_{i_1}, x_{i_2}, \cdots), \sum_{\mathbf{i}'} f'(x_{l'}, x_{i'_1}, x_{i'_2}, \cdots) \rangle \, z_{l'}^0 & (8) \\ &= \sum_{l'} \sum_{\mathbf{i}} \sum_{\mathbf{i}'} \langle f(x_l, x_{i_1}, x_{i_2}, \cdots), f'(x_{l'}, x_{i'_1}, x_{i'_2}, \cdots) \rangle \, z_{l'}^0. & (9) \end{aligned}$$

Using the definition of $\mathbf{i}$, we can merge $\mathbf{i}$ and $\mathbf{i}'$ to form a new multi-index $\mathbf{j}$ such that the equation holds for function $f''$

$$u_l^{m+1,h} = \sum_{l'} \sum_{\mathbf{j}} f''(x_l, x_{l'}, x_{j_1}, x_{j_2}, \cdots) z_{l'}^0 \tag{10}$$

Since the sum in line 9 runs through subsets of sites $\mathbf{i}$ and $\mathbf{i}'$ with size no larger than $2^m - 1$ in our assumption, the sum in Eq. 10 is over all multi-index $\mathbf{j}$ of size $\leq 2^{m+1} - 2$. Furthermore, since $z_{l'}^0$ only depends on $x_{l'}$, we can absorb the term $z_{l'}^0$ into $f''(x_l, x_{l'}, x_{j_1}, x_{j_2}, \cdots)$ to form a new function $f'''$ that depends on the same arguments as $f''$ so that

$$u_l^{m+1,h} = \sum_{l'} \sum_{\mathbf{j}} f'''(x_l, x_{l'}, x_{j_1}, x_{j_2}, \cdots). \tag{11}$$

Note that this is only possible if we use the raw embedding vector $z_l^0$ as the value vector in Eq. 6. To see why this is the case, let us assume that we use the actual output from the previous layer $z_l^m$ as the input (as in regular attention mechanisms). The $z_l^m$ vector then will contain contributions from many positions not present in the argument of the function $f''(x_l, x_{l'}, x_{j_1}, x_{j_2}, \cdots)$, so that we cannot convert the function $f''$ to a function $f'''$ with the exact same arguments.

Summing over positions $l'$, we can further simplify Eq. 11 it to

$$u_l^{m+1,h} = \sum_{\mathbf{k}:|\mathbf{k}| \leq 2^{m+1}-1} f''''(x_l, x_{k_1}, x_{k_2}, \cdots), \tag{12}$$

for some new function $f''''$. Finally, since concatenating the output of all attention heads and applying a linear transformation to this concatenated output position-wise is linear with respect to $u_l^{m+1,h}$, the final output for the $m + 1$ MHA layer can be written as

$$z_l^{m+1} = \sum_{\mathbf{k}:|\mathbf{k}| \leq 2^{m+1}-1} h(x_l, x_{k_1}, x_{k_2}, \cdots), \tag{13}$$

for some function $h$. Given the argument we made above, the sum is over all multi-indices $\mathbf{k}$ for subsets with no more than $2^{m+1} - 1$ positions. □

With this lemma, we can prove our main result.

**Proposition 1.** *The latent phenotype $\phi$ of an epistatic transformer model with $M$ MHA layers contains interactions up to order $2^M$. Specifically, $\phi$ can be expanded as*

$$\phi(x) = \sum_{k=1}^{2^M} \sum_{|\mathbf{i}|=k} e(x_{\mathbf{i}_1}, \cdots x_{\mathbf{i}_k}) + b, \tag{14}$$

*where $e(x_{\mathbf{i}_1}, \cdots x_{\mathbf{i}_k})$ is some function depending on $k$ positions indexed by $\mathbf{i}_1, \cdots \mathbf{i}_k$, and $b$ is a constant.*

*Proof.* To begin, we know that the output of the final layer of the model is

$$z_l^M = \sum_{\mathbf{k}:|\mathbf{k}| \leq 2^M - 1} f_{l,\mathbf{k}}(x_l, x_{k_1}, x_{k_2}, \cdots), \tag{15}$$

where we consider $\mathbf{k}$ to be a set and use this $f_{l,\mathbf{k}}$ notation to emphasize that the function depends on the model parameters $\theta$, the focal position $l$, and the other sites $\mathbf{k}$ $l$ has interacted with in the MHA layers.

Recall that the vectors $z_l^M$ are flattened and used to calculate a weighted sum to produce the hidden phenotype $\phi$

$$\phi(x) = \sum_l w_l^T z_l^M + b \tag{16}$$

before the final sigmoid activation. Using the expression of $z_l^M$ in Eq. 15, this becomes

$$\phi(x) = \sum_l \sum_{\mathbf{k}:|\mathbf{k}| \leq 2^M - 1} w_l^T f_{l,\mathbf{k}}(x_l, x_{k_1}, x_{k_2}, \cdots) + b. \tag{17}$$

Grouping together terms with $f$ containing the same number of arguments, we find

$$\phi(x) = \sum_{k=1}^{2^M} \left( \sum_l \sum_{\mathbf{k}:|\mathbf{k}|=k, l \in \mathbf{k}} w_l^T f_{l,\mathbf{k}}(x_l, x_{k_1}, x_{k_2}, \cdots) \right) + b. \tag{18}$$

The summand for $k$ consists of scalar-valued functions evaluated at all subsets of positions of size $k$. Therefore this equation can be expressed in the form of Eq. 14. This proves our main result. □

#### 2.3 Calculation of marginal epistasis

In this section, we provide technical details for the methods used to interpret the structure of specific epistasis in the epistatic transformer models trained on the GRB2-SH3 abundance dataset ('GRB-abundance'). Our analyses were focused on the latent phenotype $\phi$ inferred by the 3-layer epistatic transformer model, which contains only specific interactions up to order 8 and no nonspecific epistasis.

Specifically, the marginal effect of a site $l$ contributing to interaction order $k$ is defined as the proportion of variance explained by the all interactions of order $k$, involving site $l$. Mathematically, this is can be calculated as

$$V_{k,l} = \frac{\boldsymbol{\phi}^T P_{k,l} \boldsymbol{\phi}}{\boldsymbol{\phi}^T P_k \boldsymbol{\phi}}. \tag{19}$$

Here $\boldsymbol{\phi}$ is the latent phenotype inferred by the 3-layer epistatic transformer model. Like before $P_k$ is the projection matrix to the subspace spanned by $k$-way interactions. Similarly, $P_{k,l}$ is the projection to the subspace spanned by $k$-way interactions involving the focal site $i$, which can be calculated as the elementwise (Hadamard) product of two matrices

$$P_{k,l} = P_{1,l} \odot P_{k-1,-l}, \tag{20}$$

The two matrices can be defined elementwise as

$$P_{1,l}(x, x') = \begin{cases} a-1 & x_l = x'_l \\ -1 & x_l \neq x'_l, \end{cases} \tag{21}$$

and

$$P_{k-1,-l}(x,x') = W_{k-1}\left(\sum_{j\neq l} \mathbf{1}_{x=x'}\right), \tag{22}$$

where $\sum_{j\neq i} \mathbf{1}_{x=x'}$ simply calculates the Hamming distance between the two sequences while ignoring site $l$. $W_{k-1}$ is the Krawtchouk polynomial of order $k-1$, which has the same definition as Eq. 3. Intuitively, $P_{1,l}$ captures the main effects of site $l$, while $P_{k-1,-l}$ captures the interactions among all combinations of $k-1$ sites beside $l$. The product of these two matrices together captures all $k$-way interactions between site $l$ and all other sites. Note that in Eq. 19, $\boldsymbol{\phi}$ is the full size sequence-function relationship, which the predictions for all $a^L$ possible genotypes. Thus, direct calculation of the numerator and denominator in Eq. 19 is impossible for large sequence spaces. Here we rely on a mathematical result known as the kernel trick [1] to simplify the calculation.

We start by fitting a Gaussian process to the predictions made for all genotypes in the data, namely $\boldsymbol{\phi}_B$. Gaussian process regression is a Bayesian method that relies on a Gaussian prior distribution, which can be completely characterized by the kernel function $K$ specifying the covariance between any pairs of genotypes. Here we chose to parameterized $K$ as a linear sum of covariance functions, each corresponding to a prior distribution when there is only epistatic interactions among a given number of sites, which coincide with the projection matrix for the corresponding subspace $P_k$.

$$K_{\widehat{\boldsymbol{\lambda}}}(x,x') = \sum_{k=1}^{L} \widehat{\lambda}_k P_k(x,x'). \tag{23}$$

Here, $\widehat{\boldsymbol{\lambda}} = \widehat{\lambda}_1, \widehat{\lambda}_2, \cdots, \widehat{\lambda}_L$ are hyperparameters of the model, each proportional to the variance of interaction coefficients of a given order when drawing samples from the prior. Here we choose the hyperparameters by maximizing the Bayesian evidence of the data $\boldsymbol{\phi}_B$, that is

$$\widehat{\boldsymbol{\lambda}} = \text{argmax}_{\boldsymbol{\lambda}} \left[-\log|K_{\boldsymbol{\lambda}}(X_B,X_B)| - \boldsymbol{\phi}_B^T K_{\boldsymbol{\lambda}}(X_B,X_B)^{-1}\boldsymbol{\phi}_B\right] \tag{24}$$

Here $X_B$ contains all genotypes in the data, such that $K_{\boldsymbol{\lambda}}(X_B,X_B)$ contains the covariance between all genotypes in the data. Now we can infer the full sequence function relationship using standard results in Gaussian process regression

$$\boldsymbol{\phi}_{\text{GP}} = K_{\boldsymbol{\lambda}}(X,X_B)(K_{\boldsymbol{\lambda}}(X_B,X_B))^{-1}\phi_B. \tag{25}$$

$\boldsymbol{\phi}_{\text{GP}}$ contains the posterior means (maximum a posterior) estimates for all possible genotypes. To see how this simplifies our calculation, we first express $(K_{\boldsymbol{\lambda}}(X_B,X_B))^{-1}\phi_B = \boldsymbol{\omega}$, which contains weights for making predictions by combining columns of the matrix $K_{\boldsymbol{\lambda}}(X,X_B)$. We can then plug Eq. 25 into the numerator of Eq. 19 to get

$$\boldsymbol{\phi}_{\text{GP}}^T P_{k,i}\boldsymbol{\phi}_{\text{GP}} = (K_{\boldsymbol{\lambda}}(X,X_B)\boldsymbol{\omega})^T P_{k,i}K_{\boldsymbol{\lambda}}(X,X_B)\boldsymbol{\omega} \tag{26}$$

$$= \boldsymbol{\omega}^T K_{\boldsymbol{\lambda}}(X_B,X)P_{k,i}K_{\boldsymbol{\lambda}}(X,X_B)\boldsymbol{\omega} \tag{27}$$

$$\tag{28}$$

Note that the square matrix $K_{\boldsymbol{\lambda}}(X_B,X)P_{k,i}K_{\boldsymbol{\lambda}}(X,X_B)\boldsymbol{\omega}$ has the same dimension as the data. Therefore, our calculation will be greatly simplified (scaling with the number of data, instead of the size of the whole sequence space) if we can evaluate this matrix efficiently. This calculation is based on the key fact that the $a^L \times a^L$ orthonormal matrix $P_{k,i}$ spans an orthogonal subspace of $P_k$, therefore it is invariant to the projection by $P_k$, i.e. $P_kP_{k,i} = P_{k,i}$. We then re-express the matrix $K_{\boldsymbol{\lambda}}(X_B,X)P_{k,i}K_{\boldsymbol{\lambda}}(X,X_B)\boldsymbol{\omega} = A^TK_{\boldsymbol{\lambda}}(X,X)P_{k,i}K_{\boldsymbol{\lambda}}(X,X)A$, where $A = \begin{bmatrix} I_m & \mathbf{0}\end{bmatrix}^T$ is a matrix, when applied to the matrix $K_{\boldsymbol{\lambda}}(X,X)$ only preserves the columns corresponding to $B$. Putting these results together, we have

$$
\begin{aligned}
&A^T K_{\boldsymbol{\lambda}}(X,X) P_{k,i} K_{\boldsymbol{\lambda}}(X,X) A && (29)\\
=&A^T \sum_{j=1}^{L} \widehat{\lambda}_j P_j(X,X)^T P_{k,i}(X,X) \sum_{j=1}^{L} \widehat{\lambda}_j P_j(X,X) A && (30)\\
=&A^T \widehat{\lambda}_k^2 P_{k,i}(X,X) A && (31)\\
=&\widehat{\lambda}_k^2 P_{k,i}(X_B, X_B). && (32)
\end{aligned}
$$

Thus, the squared norm of the projection of the solution $\boldsymbol{\phi}_{\text{GP}}$ can be calculated exactly as

$$
\boldsymbol{\phi}_{\text{GP}}^T P_{k,i}(X,X) \boldsymbol{\phi}_{\text{GP}} = \widehat{\lambda}_k^2 \times \boldsymbol{\omega}^T P_{k,i}(X_B, X_B) \boldsymbol{\omega}, \tag{33}
$$

where $\boldsymbol{\omega}$ and $P_{k,i}(X_B, X_B)$ can be readily calculated. The denominator in Eq. 19 can be easily calculated by simply replacing $P_{k,i}$ with $P_k$, such that

$$
\boldsymbol{\phi}_{\text{GP}}^T P_k(X,X) \boldsymbol{\phi}_{\text{GP}} = \widehat{\lambda}_k^2 \times \boldsymbol{\omega}^T P_k(X_B, X_B) \boldsymbol{\omega}. \tag{34}
$$

To calculate the contribution of pure $k$th order interactions formed by a subset $S$ of $k$ positions for the sparsity analysis performed in the main text, we simply replace the projection matrix $P_k$ in Eq. 33 with the matrix for projecting to the subspace of pure $k$th order interactions, which is defined by the $k$-fold Hadmard product

$$
P_S(x, x') = \odot_{l \in S} P_{1,l}(x, x'), \tag{35}
$$

with $P_{1,l}(x, x')$ given by Eq. 21. This gives us the variance explained by $k$th-order interactions among sites in $S$. To generate their variance component, we can simply divide this by Eq.34, which is the total variance due to $k$th order interactions.

All Gaussian process modeling was performed in python package GPytorch, with 1 NVIDIA Ampere A100 GPU with 80GB memory.
